# Episodic recruitment of attractor dynamics in frontal cortex reveals distinct mechanisms for forgetting and lack of cognitive control in short-term memory

**DOI:** 10.1101/2024.02.18.579447

**Authors:** Tíffany Oña-Jodar, Genís Prat-Ortega, Chengyu Li, Josep Dalmau, Albert Compte, Jaime de la Rocha

**Author notes:** Co-senior authorship (Albert Compte, Jaime de la Rocha).

## Abstract

Short-term memory (STM) is prone to failure, especially during prolonged memory maintenance or under limited cognitive control. Despite predictive mechanistic frameworks based on persistent neural activity and attractor states, a direct assessment of network dynamics during multifactorial STM failure is still missing. We addressed this in a delayed-response task where mice maintained a prospective response during a long variable delay. Mice behavior episodically switched between a task-engaged state described by an attractor model, and a task-disengaged state purely determined by previous choices. During task engagement, the anterolateral motor cortex (ALM) showed delay persistent activity stably encoding correct choices, whereas the encoding reversed during the delay in error trials. In contrast, in task-disengaged phases ALM showed no clear traces of attractor dynamics and instead exhibited enhanced synchrony at ∼ 4-5Hz. Thus, ALM switches between distinct error-generating dynamics: in control-capable trials, transitions between memory attractors cause forgetting errors, whereas non-memory errors are caused by the dissociation of ALM during the mnemonic period reflecting the lack of cognitive control.

## INTRODUCTION

Short-term memory (STM), the ability to retain information in the brain during short periods of time, is crucial as a component of working memory in multiple cognitive functions such as planning, abstract thought, language, and decision making (Baddeley 2003). In spite of its versatility and rapid dynamics, STM is markedly limited both in its storage capacity (Halford, Cowan, and Andrews 2007) and its short-lived nature as it suffers degradation through time at various time scales: from rapid mnemonic time scales of seconds (Mercer and Barker 2020) to history-based interference (Fritsche, Mostert, and de Lange 2017), time-on-task effects (Krimsky et al. 2017), and mind wandering (Mrazek et al. 2012). Understanding the mechanistic basis of these limits seems imperative as they are implicated in multiple brain disorders such as schizophrenia or anti-NMDA-receptor encephalitis (Johnson et al. 2013; Guasp et al. 2022).

Electrophysiology studies have extensively described memory-related neural states during STM tasks in monkeys (Funahashi, Bruce, and Goldman-Rakic 1989; Fuster and Alexander 1971; Kubota and Niki 1971; E. K. Miller, Erickson, and Desimone 1996; Romo et al. 1999; Miyashita and Chang 1988; David J. Freedman and Assad 2006; D. J. Freedman et al. 2001) and rodents (Baeg et al. 2003; Fujisawa et al. 2008; Erlich, Bialek, and Brody 2011; Harvey, Coen, and Tank 2012; Inagaki et al. 2018, 2019; Goard et al. 2016; Liu et al. 2014). Many experiments have found neurons in associative cortices that, during the delay, exhibit selective persistent activity encoding the retrospective memory of the stimulus properties and/or the prospective response (Miyashita and Chang 1988; Mendoza-Halliday, Torres, and Martinez-Trujillo 2014; Liu et al. 2014) (Inagaki et al. 2019; Funahashi, Bruce, and Goldman-Rakic 1989). These neural dynamics can be conceptualized in a mechanistic framework that describes memory states as equilibrium states, or attractors, of a neural network. In the attractor state a small, specific, neural ensemble exhibits sustained, elevated (or decreased) activity that is maintained by positive feedback loops (Amarimber 1972; Wilson and Cowan 1972; Amit and Brunel 1997). However, the attractor framework is challenged by evidence showing that delay activity can exhibit complex temporal dynamics that appear to be at odds with equilibrium states (Lundqvist, Herman, and Miller 2018). One example are STM navigation tasks in rodents, where the encoding seems to consistently occur in sequences during which neurons fire transiently (Harvey, Coen, and Tank 2012; Fujisawa et al. 2008; Zhu et al. 2020) and for which alternative models based on sequential neural activations have been proposed (Rajan, Harvey, and Tank 2016; Goldman 2009; Laje and Buonomano 2013).

Because failures of STM can greatly constrain existing models, memory errors have received substantial experimental attention (Yang et al. 2022; Inagaki et al. 2019; Rossi-Pool et al. 2017; Funahashi, Bruce, and Goldman-Rakic 1989; Sakai and Miyashita 1991; David J. Freedman and Assad 2006; Vergara et al. 2016). Most studies found that persistent activity in error trials is decreased or absent (Sakai and Miyashita 1991; David J. Freedman and Assad 2006; Funahashi, Bruce, and Goldman-Rakic 1989; Vergara et al. 2016) whereas others found delay activity robustly encoding the incorrect upcoming choice throughout the delay period (Yang et al. 2022; Inagaki et al. 2019). Independently of the observed effect, these results still lack a common mechanistic explanation that describes memory failure as a result of altered network dynamics. Indeed, testing model predictions against experimental data in error trials is difficult. This is in part due to some tasks being too simple and mnemonic periods too short for subjects to make a sufficient number of errors (see e.g. (Funahashi, Bruce, and Goldman-Rakic 1989; Sakai and Miyashita 1991)), but also because behavioral errors are mechanistically heterogeneous and pinpointing the cause of a specific error is difficult. In general, memory errors can be due to defective memory loading, maintenance or retrieval (Mayer and Park 2012; Bays et al. 2011;

Jonides et al. 2008; Alleman et al. 2023) and there can be errors unrelated to the task contingency, such as lapses (Adam et al. 2015; deBettencourt et al. 2019). The use of STM during long periods requires sustained cognitive control (Kahneman 2011), the lack of which may also cause decision errors seemingly associated with a lack of memory usage rather than a failure of memory *per se* (Sawagashira and Tanaka 2021). In a similar vein, recent work has shown that mice performing decision making tasks can switch between behavioral strategies, corresponding to a different level of task engagement, during the session (Ashwood et al. 2022; Bolkan et al. 2022), further challenging the analysis of error choices.

One specific type of STM error has been shown to be consistent with attractor dynamics in the prefrontal cortex. In a visuo-spatial memory-guided task in monkeys (Funahashi, Bruce, and Goldman-Rakic 1989; Constantinidis and Goldman-Rakic 2002), small trial-by-trial variations in the delay persistent activity of prefrontal cortex neurons predicted small deviations in the delayed response of the subjects (Wimmer et al. 2014). This correlation was a prediction of a continuous attractor model operating in the degenerate regime under the influence of noise: activity fluctuations during the delay period make the memory *bump* randomly move across a continuum of states, an effect that increases with the delay duration (Wimmer et al. 2014; Burak and Fiete 2012; Compte et al. 2000). This theoretical understanding of the mechanisms of errors in this STM task is difficult to test causally in primates, and a parallel approach in rodents is still lacking. Interestingly, the circuit mechanisms of simpler categorical STM tasks in rodents have been identified in multiple brain areas and seem consistent with discrete attractor models (Inagaki et al. 2019; Kopec et al. 2015). It remains to be shown whether the maintenance errors that these models predict are consistent with the circuit dynamics observed during memory error trials (Erlich, Bialek, and Brody 2011; Inagaki et al. 2019)

Here, we investigate the neural mechanisms that can make STM fail in mice trained to perform a two-alternative delayed-response task with long variable delays (i.e. up to ten seconds). We analyze different error types, with the aim of specifically isolating and characterizing memory maintenance errors, i.e. those responsible for forgetting. We develop a two-state Hidden Markov model to describe the within-session episodic switching between task-engaged and task-disengaged phases. We then leverage this model to analyze the encoding of the prospective choice in populations of ALM neurons during these two phases. During cognitive engagement phases, population dynamics in long-delay error trials have the predicted features of forgetting errors in an attractor model operating in the winner-take-all regime: correct response loading upon stimulus presentation, and transition from the correct to the incorrect memory attractor during the delay. In task-disengaged trials, the same ALM circuit exhibits very different dynamics with a lack of clear sustained memory activity and no signatures of memory maintenance errors. Our results not only support attractor network dynamics as the mechanism underlying STM but extend this framework by showing how cognitive engagement modulates network dynamics as memory circuits in the cortex transition between control-capable and control-limited states.

## RESULTS

### Intermittent engagement in short-term memory

To study the mechanisms of STM maintenance and the sources of errors during this period, we trained 48 mice to perform an auditory two-alternative delayed choice (2ADC) task (Fig 1a). Animals were trained to listen to a sound stimulus that was presented laterally and lick the corresponding lickspout. If animals licked the port on the side of the presented sound, they received a water reward. Between sound presentation and response, a delay was imposed, which could randomly take durations of 0.1, 1, 3 or 10 s. After the delay, a motor brought the water spouts within reach of the animal, functioning as the Go cue. We found that the choice accuracy, i.e. the fraction of correct responses, decayed with delay duration, implying that at least a fraction of the errors were caused by memory degradation during the mnemonic period (Fig. 1b; one-way ANOVA: *f* = 12.791, *p* = 2.2 × 10^−7^). Each animal’s curve was parametrized by the regression slope (mean ± SD = 0.009 ± 0.004 s^-1^) and the lapse rate, defined as one minus the accuracy at zero delay (0.15 ± 0.055). Inter-individual variability in overall accuracy (i.e. averaged accuracy across delays) did not correlate with the slope (Fig. 1d; R^2^ = 0.067, *p* = 0.14) but it correlated strongly with the lapse rate (Fig. 1e; R^2^ = 0.881, p<0.001). Therefore, the main source of inter-individual variability in accuracy did not come from differences in the stability of memory across time but rather from differences in lapses, supposedly caused by some factor other than memory maintenance. We found that animals showed a consistent tendency towards repeating the previous choice despite the absence of repeating structure in the presented stimuli. We quantified this tendency using the Repeating Bias index (RB) defined as the corrected probability to repeat the previous choice (0.56 ± 0.04; Methods). Critically, RB was not affected by delay length (Fig. 1f and g ; one-way ANOVA; *f* = 0.301, *p* = 0.82), indicating that choice repetition was not a default strategy that subjects applied for instance when forgetting during the delay or when losing motivation specifically in long-delay trials. When we correlated RB with lapse rate, we observed a clear positive correlation, with higher lapse rates found in animals with higher RB (Fig. 1h; R^2^= 0.192, p<0.008). Moreover, repeated and correct responses did not occur in an uncorrelated manner along the trial sequence but they tended to cluster in short episodes of 5-20 trials, which we call session phases, as shown by autocorrelation analysis (Fig. 1i-j).

**Figure 1.**
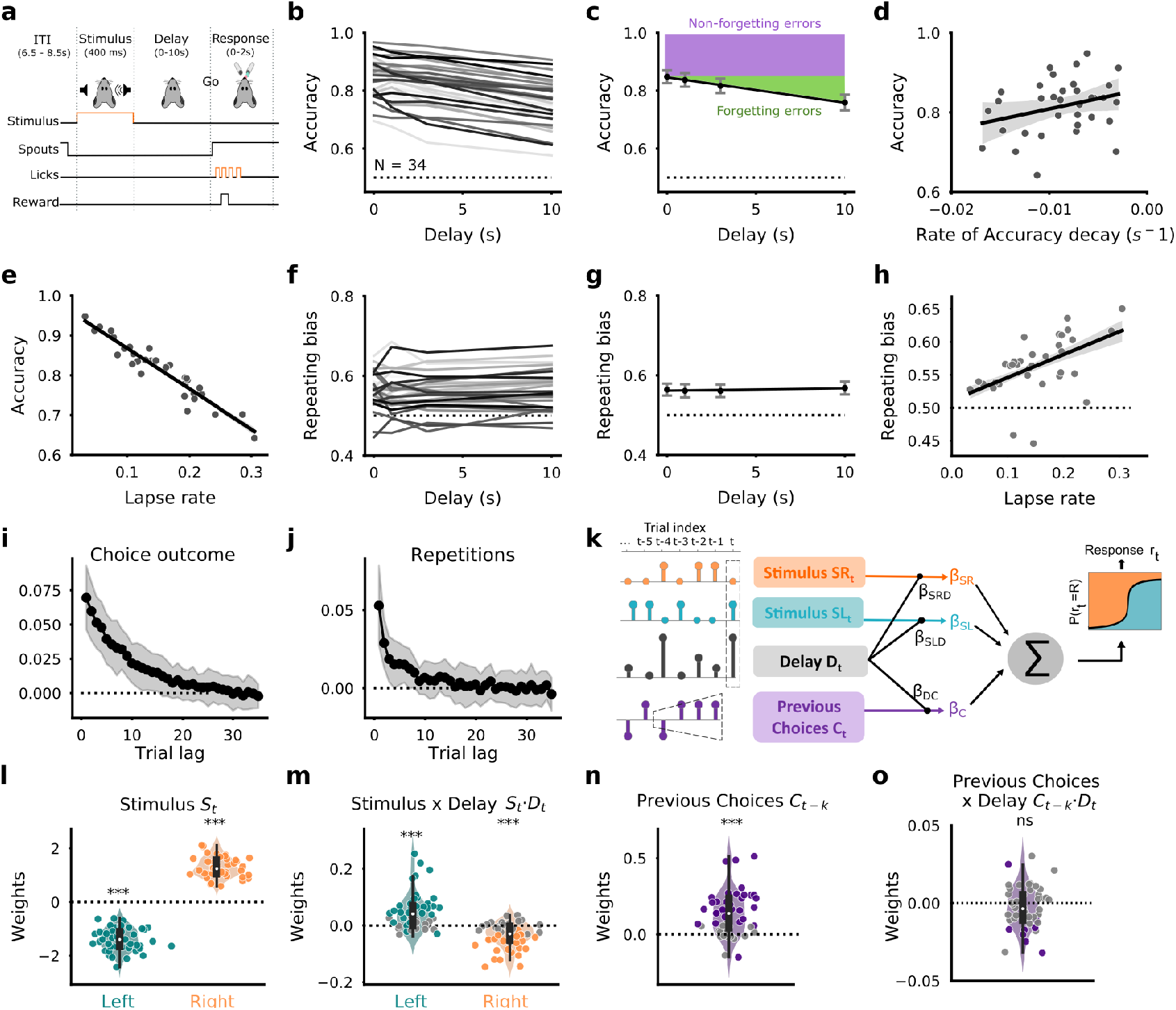
Behavioral quantification in 2ADC task using variable delays. **a**, Sketch of the task: a stimulus coming from either a Left (L) or a Right (R) speaker is presented (duration 400 ms). Following a variable delay, two ports are approached to the animal, which has then to lick the corresponding side (Left Stimulus → Left port and Right Stimulus → Right port). Correct responses are rewarded with 2.5 μl of water, and incorrect ones are followed by a dim light and a 2 s timeout. Retraction of the ports occurs 2.5 s before the next stimulus onset. **b-c**, Choice accuracy versus delay duration. Individual animals (curves in b; Group #2, n=34) and mean and s.e.m. across animals (dots in c). Regression slope was -0.009 s^-1^ (line in c). Colored areas represent the average fraction of forgetting (green) and non-forgetting errors (purple). **d-e**, Delay-averaged accuracy versus rate of accuracy decay (d) or versus lapse rate (e). Each dot represents one animal and the line represents a linear regression. **f-g**, Repeating bias (RB) versus delay duration. Individual animals (f) and mean across animals (g) with a linear regression (slope 10^−4^ s^-1^, p= 0.26). **h**, Repeating bias versus lapse rate. **i-j**, Autocorrelogram for sequences of choice outcomes (i: correct =1 vs. incorrect =0) or repetition sequence (j; repeat =1 vs alternate =0). Autocorrelograms were corrected to remove the influence of within-session drifts (see Methods). Dots indicate mean across animals and the shaded area represents the 95% C.I. **k**, Schematic of the GLMM fitted to the choices of each individual animal. Regressors include the stimulus (separated into Right and Left stimuli), a weighted sum of previous responses, the interaction of delay with these and session as a random factor (Methods). **l-o**, fitted GLMM weights for the stimulus (l), the interaction between stimulus and delay (m), the action trace *C*_*t*_ accounting for previous response (n) and its interaction with delay (o). Dots represent individual subjects (Groups #1 and #2, n=48). Colored dots indicate individually significant weights (*p* < 0.05).

We fitted a generalized linear mixed model (GLMM) to the responses of each animal to quantify the simultaneous impact of the current stimulus and previous responses and their interaction with the current delay (Fig. 1k). The analysis confirmed that delay interacted with the stimuli by decreasing their impact (Fig. 1l-m) but the influence of previous choices (Fig. 1n and Supplementary Fig. 1) did not depend significantly on the delay length (Fig. 1o). Average accuracy decreased towards the end of the session, a pattern commonly observed in trial-based tasks as animals get satiated. This trial-dependent decrease was accompanied with an increase in RB but with no change in memory degradation rate (Supplementary Fig. 2). Together, these results show that, during the task, mice make memory errors and repeating lapses, with rates that are preserved or vary substantially across conditions (i.e. early vs late in the session), respectively. In addition, repeating lapses, which can explain individual differences in task performance (Fig. 1e), appear decoupled from delay-sensitive memory maintenance.

### The repeating behavior cannot be reproduced by the attractor model

We next aimed to extend the double-well attractor model (DW) to account for both the decrease of accuracy with delay as well as the delay-independent RB. The DW model, previously used to capture behavior and population activity in 2ADC tasks (Finkelstein et al. 2021b; Inagaki et al. 2019), is analytically tractable while preserving a mechanistic network foundation (Roxin and Ledberg 2008; Wong and Wang 2006; Prat-Ortega et al. 2021) and can be parametrized with four parameters: the probabilities *P*_*L*_ and *P*_*R*_ to load the correct memory upon presentation of the Left or Right stimulus, respectively; the height of the barrier *α* which, for a given noise magnitude, set the transition rate between the two wells during the delay period (Prat-Ortega et al. 2021); and the parameter *μ* which introduced an asymmetry in the depth of the two wells. To bias the DW towards previous choices we incorporated a linear dependence on the trial-by-trial *action trace C*_*t*_, defined as the weighted sum of previous responses (Fig. 2a), in either (1) the initial loading probabilities *P*_*L*_ and *P*_*R*_ (*DW repeat-biased loading*, DW-L), (2) the asymmetry parameter *μ* (*DW repeat-biased maintenance*, DW-M), or (3) in both parameters (*DW repeat-biased both*, DW-B; Fig. 2a; Methods). Using Kramers’ reaction-rate theory (Kramers 1940), we obtained approximate analytical expressions for the models’ likelihood to generate the two output choices allowing us to fit each of the DW model extensions, including the classic *repeat-independent* DW (*DW*) with no dependence on *C*_*t*_, to the choices of each mouse (Methods). All fitted models quantitatively reproduced the decrease in accuracy with delay, reflecting the occurrence of noise-driven transitions between the two model attractors (Fig. 2b). However, all DW models failed to generate an above-chance RB that was constant across delays (Fig. 2c): when the memory loading was biased towards previous responses (DW-L), the RB decreased with delay as the probability to observe a transition increased (Fig. 2c). In contrast, when the attractor corresponding with previous choices was made deeper (DW-M), the RB increased with delay, starting at chance for *D*=0, as the probability to fall into the deeper well increased (Fig. 2c). Combining the two mechanisms (DW-B) improved the fit but the model still exhibited a significant dependence of the RB on delay (Fig. 2c). Thus, biasing choices using any of the DW parameters inevitably induced a dependence on the delay length, demonstrating that this model alone could not reproduce this important qualitative aspect of the animals’ behavior.

**Figure 2.**
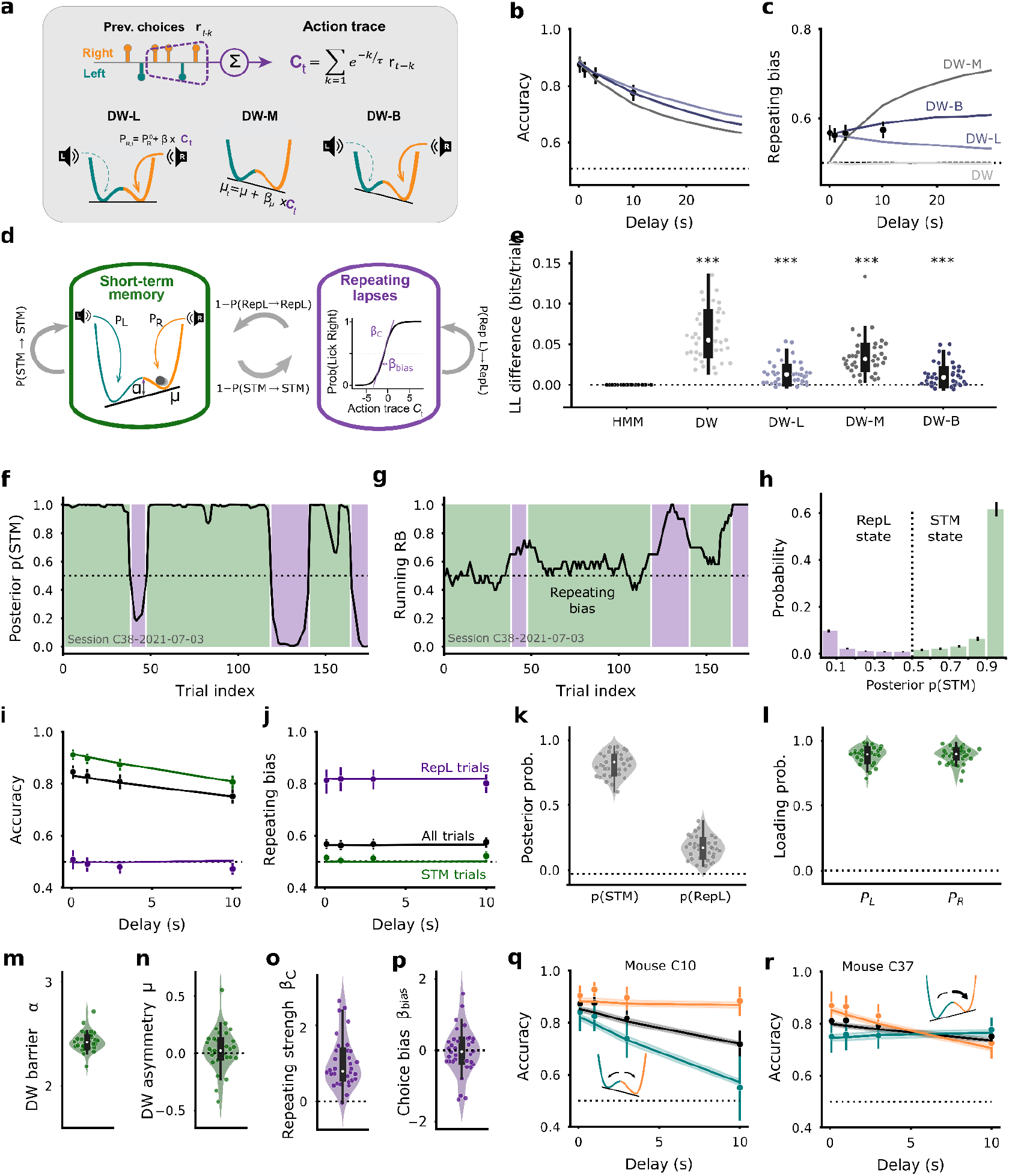
A two-state Hidden Markov Model with Double Well dynamics can fit mouse behavior. **a**, Schematics of three extensions of the DW attractor model that incorporate a repeating bias. Action trace *C*_*t*_ in each trial, defined as the weighted sum of previous choices (top), impacts either the memory loading probabilities (DW-L), the asymmetry of the DW during memory maintenance (DW-M) or both parameters (DW-B). **b-c**, Fits of the accuracy and RB versus delay by each DW model (lines). Delay range was increased to 30 s to better illustrate the qualitative dependence. Dots represent the mean ± sem of the data (Group #2, n=34 mice). **d**, Schematics of the HMM model defined by 6+3 parameters defining behavior within each state, (*P*_*L*_, *P*_*R*_, *α, μ, β*_*c*_, *β*_*0*_), and the probabilities p(STM→STM), p(RepL→RepL) (state transitions) and Π (probability to start the session in STM, not shown in the schematic; see Methods). **e**, Cross-validated log-likelihood difference per trial between the different DW extensions and the HMM. Dots represent individual mice (asterisks represent paired-test against HMM values). **f**, Posterior probability p(STM) of being in the STM state from an example session. Shaded areas represent STM (green) and RepL (purple) session phases classified using the 0.5 probability threshold (dotted). **g**, RB trace computed using a 20-trial running window for the session shown in f. **h**, Histogram of the HMM posterior probability p(STM) averaged across all animals. **i-j**, Accuracy and RB versus delay for all trials (black), STM phase trials (green) and RepL phase trials (purple). Dots represent the mean ± sem of the data and the lines the model fits. **k**, Mean posterior probabilities p(STM) and p(RepL) provided by the fitted HMM for each animal/session. **l-p** HMM-fitted values for the memory loading probabilities *P*_*L*_ and *P*_*R*_ (l), DW barrier height *α* (m), asymmetry parameter *μ* (n), sensitivity to the action trace β _*c*_ (o) and choice bias β _*0*_ within the RepL state (p). Other fitted parameters of the HMM are shown in Supplementary Fig. 3a. **q-r**, Accuracy vs delay for two individual animals exhibiting different idiosyncratic delay-dependent choice biases (dots) and corresponding HMM fits (lines). Accuracy was computed using all (black), Right-stimulus (orange) or Left-stimulus trials (cyan). Example mice show a delay-increasing lateral bias (q) or crossing-pattern with opposite side preference at short and long delays (r). Inset shows the parameterization of the DW used by the HMM to capture each delay dependence of the choice bias. Averages and individual mouse dots in panels b-c, e, h-p come from Group #2 (n=34 mice) whereas in panels k-p come from Groups #1 and #2 (n=48).

### Behavior can be reproduced using a Hidden Markov Model that includes attractor dynamics

We reasoned that the observed decoupling of repeating lapses from the dynamics of memory maintenance was probably reflecting two separate behavioral strategies that animals used in different trials to engage in the task, one guided by the stimulus and another one solely based on previous actions. To formalize this idea, we built a Hidden Markov Model (HMM) with two behavioral states representing each of these strategies: in the STM state (STM), animal choices were described by the dynamics of a repeat-independent DW model, whereas in the Repeating Lapses state (RepL), choices ignored the stimulus and were only guided by the action trace *C*_*t*_ (Fig. 2d; Methods). According to this model, accuracy under the RepL state is 0.5 whereas under the STM state it is determined by the DW parameters describing memory loading (*P*_*L*_ and *P*_*R*_) and maintenance (*α* and *μ*). We fitted the HMM for each mouse and compared it with the DW model extensions: the HMM explained the data better than any of the DW models in almost every mouse (Fig. 2e; number of mice not favoring the HMM were 0, 3, 0 and 6 out of 34 mice; paired t-tests, p = 2.3×10^−13^, 8.9×10^−7^, 4.5×10^−10^ and 3.2×10^−5^, for the DW, DW-L, DW-M and DW-B, respectively). Using the fitted HMM, we computed the posterior probability p(STM) of a mouse being in the STM state in each trial. For each session, we identified STM and RepL phases depending on whether p(STM) was above or below 0.5, respectively (Fig. 2f, shaded areas). Inspection of a running estimate of RB across the session confirmed that STM and RepL phases tended to exhibit low and high RB values, as expected from the model’s design (Fig. 2g). The distribution of p(STM) was strongly bimodal indicating that the model’s classification of most trials was unambiguous (Fig. 2h). Importantly, the HMM could quantitatively reproduce the decrease in accuracy with delay, produced by the DW dynamics underlying the STM state, and the constant repeating bias, reflecting the choices generated from the RepL state (Fig. 2i-j). Hence, when conditioned on RepL phases, the mice accuracy was around chance and there was a substantial increase in the RB, in close agreement with the model (Fig. 2i-j, purple). Furthermore, when conditioned on STM phases, accuracy increased preserving the decay with delay whereas the RB decreased to chance, also in close agreement with the HMM (Fig. 2i-j, green). The model revealed similar behavioral structure in all mice: Mice spent 81.6 ± 11.2% of the trials in the STM state and only 18.3 ± 11.2% in the RepL state (Fig. 2k). Also, HMM-fitted parameters were quite consistent across subjects (Fig. 2l-p). The probabilities *P*_*L*_ and *P*_*R*_ for correctly loading the correct memory into the DW were 90.4 ± 6% and 90.1 ± 5.6% indicating that, according to the HMM, in approximately 10% of trials in the STM phase, mice made errors that were not caused by a failure in memory maintenance (Fig. 2l). The HMM quantitatively captured other features of the behavior such as the cumulative impact of previous responses on accuracy (Supplementary Fig. 3b), the artifactual dependence of accuracy and RB on the previous trial outcome (Supplementary Fig. 3d-e), as well as the across-trial correlation structure of outcomes and repetitions (Supplementary Fig. 3f-g). Lastly, the flexibility of the model in generating asymmetric DWs was able to capture idiosyncratic history-independent choice biases and their delay dependence in each individual animal (Fig. 2q-r and Supplementary Fig. 3c). In summary, the fitted HMM suggests that mice constantly alternate between two behavioral strategies: one requiring memory maintenance and described by a DW attractor model with negligible trial-history effects; and a possibly less effortful one, in which no stimulus processing is required, and where choices are simply guided by the inertia of previous actions. Interestingly, the model infers on a trial-by-trial basis whether mice are using one strategy or another, a powerful tool that we exploited in our subsequent analyses.

### Population activity in ALM encodes choice during the delay in STM trials

We wondered if, by leveraging the fitted HMMs, we could identify distinct neural signatures associated with memory maintenance errors in STM phases and repeating lapses during RepL phases. To this end, we recorded population neuronal activity using silicon probes in area ALM (Group #3, n=8 mice; n=40 sessions; 2,089 units, Methods), a frontal area previously associated with attractor dynamics in similar tasks (Inagaki et al. 2019; Finkelstein et al. 2021a). Simultaneously recorded neurons showed great heterogeneity in their responses across trial epochs, i.e. stimulus, delay and response (Fig. 3a). Across trials, individual neurons showed previously reported selectivity between Left and Right trials (Inagaki et al. 2019, 2018), particularly during the stimulus and choice periods (Supplementary Fig. 4). We aimed to study the neural dynamics underlying the various identified behaviors in a systematic way. So, we first focused on the population dynamics in correct trials during identified STM phases. Decoders trained to predict the current trial’s choice based on the activity of simultaneously recorded neurons at different times within the trial revealed a specific pattern of generalization to other trial periods (Fig. 3c). First, a transient Stimulus code appeared upon sound presentation, which largely decayed during the delay (Fig. 3d). Second, a memory code emerged in the delay, which generalized almost perfectly for the whole delay duration, even when training only in the last 250 ms of the delay (Delay code, Fig. 3e). Consistent with this stable coding of the upcoming response during the delay, we found neurons with remarkably steady selective average firing rate throughout the delay period (Fig. 3b). Lastly, a Response code was revealed in the response window, which was transient (Fig. 3f), appeared more dynamic and was largely orthogonal to the Delay code given the total absence of cross-generalization between the two (Fig. 3c). The same results were obtained in 10s-delay trials (Supplementary Fig. 5). The relation between the different codes was quantified using the cosine of the angle between the decoding vectors (Fig. 3g): while Delay codes trained at the beginning or at the end of the delay were almost collinear (mean cos(Early, Late) = 0.463 ± 0.317 was indistinguishable from cos(Late, Late’) = 0.537 ± 0.174, t = -0.599, p=0.55), Stimulus code and especially Response code approached orthogonality with respect to Delay code (cos(Delay, Stim) = 0.214 ± 0.339, t = 4.0, p<10^−3^; cos(Delay, Resp) = 0.085 ± 0.376, t = 1.43, p = 0.16). This suggests that different combinations of ALM neurons underlie each of these three population codes (Yang et al. 2022; Inagaki et al. 2022).

**Figure 3.**
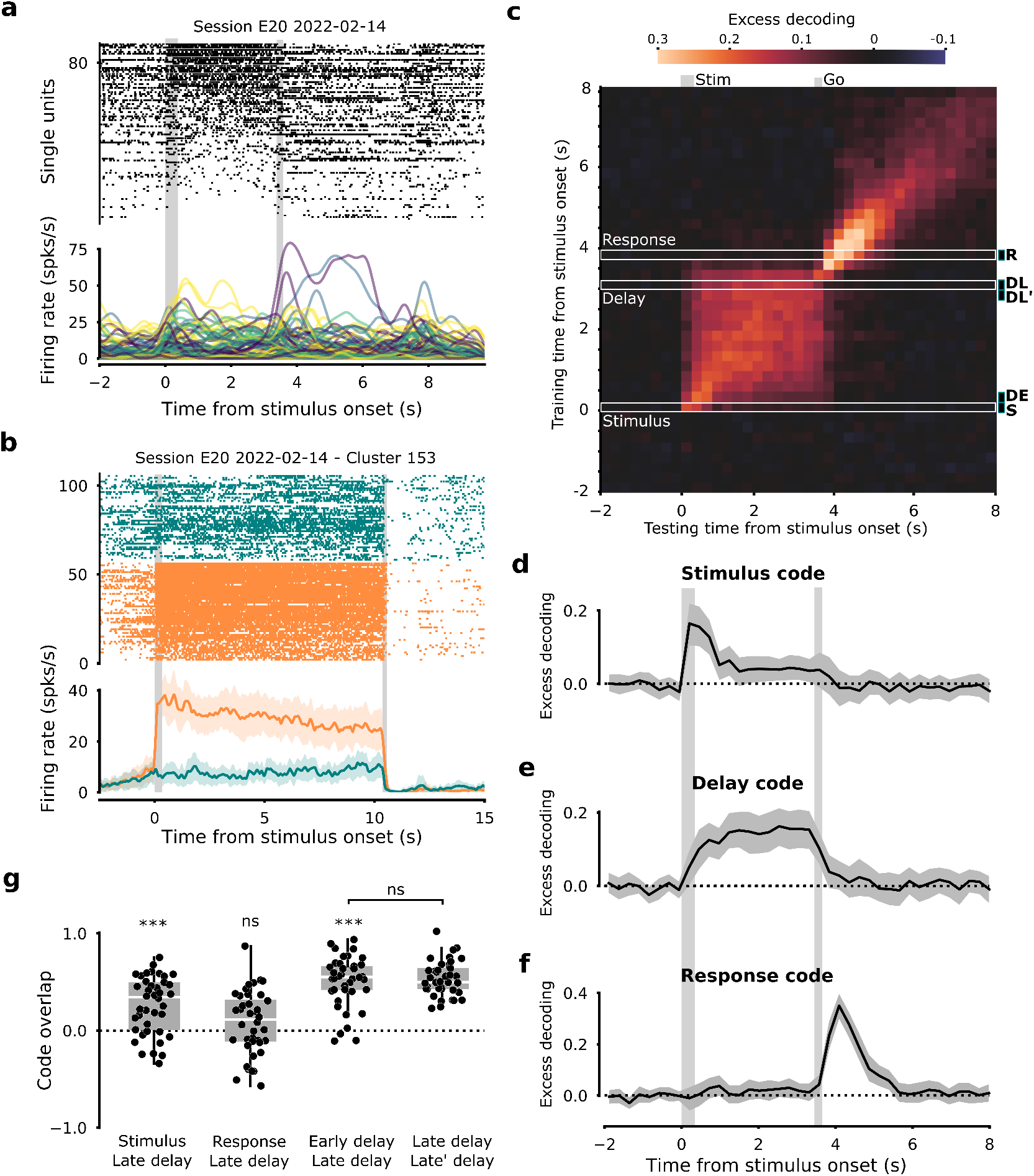
ALM spiking activity in correct STM trials reveals three different population codes. **a**, Instantaneous population activity from an example trial with a 3s delay (Mouse E20). Top raster shows spike times from 98 simultaneously recorded units, sorted according to their firing rate during the delay. Bottom plot shows instantaneous rates for each unit. Stimulus and Go cue are marked with gray rectangular shades. **b**, Post-stimulus time histograms (PSTHs) obtained from correct STM 10s-delay trials from an example neuron that exhibits selective sustained activity during the delay (session E20_20220214 cluster 153). Color indicates trial type (Right stim. orange; Left stim. blue). **c**, Average cross-validated excess decoding accuracy (Methods) from a linear decoder trained and tested using the population activity at different trial times to predict the animal’s choice. Only 3s-delay correct trials classified as STM by the HMM were used for training and testing (Group #3 with n=8 mice; n = 40 sessions; see also Supplementary Fig. 5 for 10s-delay trials). The Stimulus code, Delay code and Response code are defined by the training time intervals (0, 0.25) s, (3, 3.25) s and (3.75, 4) s (white rectangles). **d-f**, Time evolution of the excess decoding accuracy of the Stimulus, Delay and Response codes. Lines represent the average across sessions and bands are 95% C.I. **g**, Overlap between codes quantified by the cosine of the angle between pairs of decoder weight vectors trained in indicated task periods (using 3s-delay and 10s-delay trials, see Methods). The time windows used for training are indicated as vertical ticks in panel c: Delay Early (DE) was (0.5, 0.75) s, Delay Late (DL) was (3, 3.25) or (10, 10.25) s depending on the trial’s delay D and Delay Late’ (DL’) was the same as Delay Late but advanced 0.25 s. Stimulus (S) and Response (R) codes were as defined above. Dots indicate individual sessions. All plots are temporally aligned to stimulus onset.

### Population activity shows traces of memory state transitions during STM error trials

We next used these three codes, identified using correct trials in the STM phase, to interrogate our data about the mechanisms underlying erroneous responses in the STM phase. In such incorrect trials, the accuracy of the Stimulus code increased transiently during stimulus presentation, similarly to correct trials (Fig. 4a). Notice that, because in incorrect trials the stimulus and the response differ, the matching traces demonstrate that the Stimulus code represented the stimulus robustly also in incorrect trials. Thus, errors in the STM phase were not generally caused by stimulus information not reaching ALM correctly. The Response code also showed comparable decoding dynamics with respect to correct trials (Fig. 4b), despite not reaching the same peak accuracy possibly due to the decrease in the number of licks (in non-rewarded responses mice licked 1-2 times whereas in rewarded responses they made 10-15 licks). Lastly, the accuracy of the Delay code in predicting the animal’s incorrect choice showed a striking difference in long-delay trials: early in the delay, accuracy was negative, meaning that the population represented the opposite choice to the one finally executed (Fig. 4c). As the delay progressed, the initially negative encoding gradually reversed to reach positive values, comparable to the encoding observed in correct trials (Fig. 4c-d). This means that, in these error trials occurring in the STM phase, the neurons representing the prospective response during the delay switched from encoding the correct choice after stimulus onset, to encoding the opposite incorrect choice by the end of the delay. This reversal pattern could be observed on single sessions (Fig 5e), in the average response of single neurons (Fig. 4f) or in the decoding dynamics of single-trials (Fig. 4g-h). This code reversal in error trials is in fact a prediction of DW attractor models, where random fluctuations during the attractor state can cause an attractor transition from the correctly encoded memory to the alternative, erroneous choice (Supplementary Fig. 7a)(Moreno-Bote, Rinzel, and Rubin 2007; Prat-Ortega et al. 2021). Importantly, code reversals were absent when analyzing short delay trials (1 and 3 seconds; Supplementary Fig. 6), a feature also predicted by the DW: because the transition rate is so small, only for long delays (10 s) the proportion of transition errors among the total number of errors is sufficiently large for the average decoding dynamics to show a reversing trace (Supplementary Fig. 7). In total, our analysis shows that in the STM phase and specifically in long-delay trials, when animals are more prone to maintenance errors, neural populations showed dynamics consistent with transitions between memory states.

**Figure 4.**
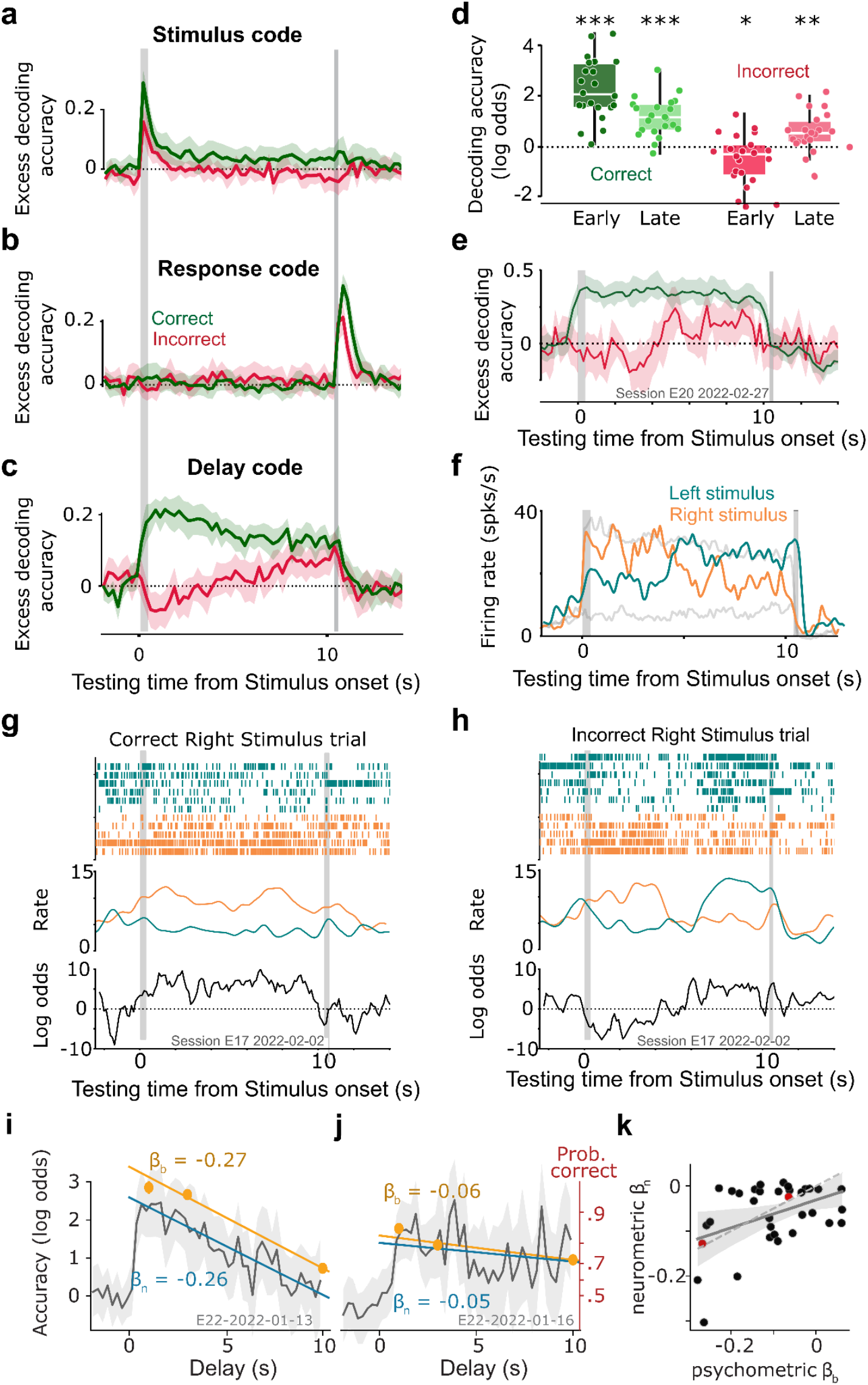
Comparison of neural activity between correct and incorrect STM trials. **a-c**, Average cross-validated excess of decoding accuracy of the Stimulus code (a), Response code (b) and Delay code (c) trained using 5-fold cross validation with all STM correct trials and tested in both STM correct (green) and incorrect (red) trials with 10s delay (n = 21, 22 and 30 sessions respectively; see Table 1 in Methods; see Supplementary Fig. 6 for 1s and 3s-delay trials). Positive excess decoding accuracy indicates proper stimulus decoding for the Stimulus code and proper response decoding for the Delay and Response codes. **d**, Log odds obtained from the Delay code averaged over the first and last second of the 10s delay (early vs. late) for both STM correct and incorrect trials. Dots indicate individual sessions. Linear mixed model on individual trials: Correct/Error, weight= 2.84, p < 10^−15^; Early/Late, weight= 1.08, p = 0.82 and interaction weight=-2.17, p < 10^−15^. Asterisks represent post-hoc Wilcoxon tests against zero (p<10^−4^; p<10^−4^; p=0.016; p=0.001, from left to right) and for early vs. late in STM correct (wilcoxon, p < 10^−4^) and STM incorrect phases (wilcoxon, p < 10^−4^). **e**, Same as in c but for an example single session. **f**, Single neuron example (same neuron as Fig 4b) firing rates for incorrect Left (2) and Right stimulus (5) trials classified as STM. Gray underlying curves depict firing rate in correct STM trials for comparison. **g-h**, Examples of a correct (g) and an incorrect (h) single trial population activity from neurons that participate in the Delay code showing selectivity for either Left (blue, n = 6) or Right (orange, n = 5) choices. Top: population spike raster grams. Middle: averaged instantaneous firing rate from each group of neurons. Bottom: instantaneous readout from the Delay code. Negative (positive) values correspond to higher probability for the Left (Right) response. The error trial (h) shows a putative state transition from the Left to the Right memory attractor around t=5s. **i-j**, for two example sessions, psychometric accuracy (orange), i.e. probability that mice make a correct response, and neurometric accuracy (gray), i.e. probability to read out the *correct* response (the one matching the stimulus) from the Delay code as a function of delay. Straight lines are the corresponding linear fits and ***β*** _b_ and ***β*** _n_ are their slopes defined as psychometric and neurometric forgetting rates. Both panels i-j share the y-axis in panel i (log odds units) and the red axis on panel j (probability units). **k**, relationship between the psychometric ***β*** _b_ and neurometric ***β*** _n_ forgetting rates obtained across sessions. Each dot represents a session and the solid line represents a linear fit. Dashed line shows the unity line. Red dots mark the two sessions shown in panels i-j.

**Table 1.** Table showing the organization of the mice into different groups depending on the experiment they were used for. Notice that some mice belong to more than one group.

| Animal groups | No. of mice | No. sessions (mean±std; range) | No. trials per session per mouse (mean ± std; range) | Total No. trials |
| --- | --- | --- | --- | --- |
| Group #1: Behavioral task with delays D = 0, 1 and 3 s.<br>Mice: C12, C15, C18, C22, C28, C34, C39, N09, N11, N22, N21, E06, E21, E18 | 14 | 22.57 ± 8.86;<br>11-39 | 355.99 ± 54.73;<br>271.18-471.25 | 109310 |
| Group #2: Behavioral task with delays D = 0, 1, 3 and 10 s.<br>Mice: C10b, C13, C19, C36-C38, N02-N05, N07, N08, N09, N24-N28, E03-E08, E10-E17, E19, E20, E22 | 34 | 21.85 ± 13.35;<br>5-51 | 358.61 ± 65.48;<br>210.6-461.23 | 232487 |
| Group #3: Electrophysiological ALM recordings<br>Mice: E04, E14, E11, E13, E17, E19, E20, E22 | 8 | 5 ± 1.87;<br>2-8 | 351.3 ± 127.38<br>88-651 | 14052 |
| TOTAL | 48 |  |  | 355849 |

We designed an analysis to test empirically the predicted association between the delay code reversals in ALM population activity and random transitions between memory states, which induce delay-dependent memory errors in DW attractor models. If transitions between memory states underlie the population dynamics during the delay, then the reversal rate of delay code should correlate across sessions with the behavioral forgetting rate in STM phases. We took all correct and incorrect STM trials together and interrogated the ALM Delay code to provide a Left vs Right response at each time point during the delay. We used these readouts to compute, for each session, a neurometric curve defined as the probability of the decoded choice to be correct, i.e. to match the current stimulus, as a function of elapsed delay time (Fig. 4i-j, gray). The decrease rate of this curve, defined as the neurometric forgetting rate, captured the reversal rate of delay neurons. Then, we compared for each session the neurometric forgetting rate with the behavioral forgetting rate, defined as the decrease rate of the psychometric curve (Fig. 4i-j orange and Supplementary Fig. 8). As predicted, the two rates were significantly correlated across sessions (Pearson r = 0.43, p = 0.0052; Spearman r = 0.32, p = 0.033; Methods; Fig. 4k). This result suggests that, during STM phases, the temporal instability of choice encoding in ALM neural activity during the delay underlies behavioral forgetting.

### Population activity shows disrupted memory representations in RepL trials

The attractor dynamics that supported choice maintenance in the STM phase were, however, very different during the RepL phase. Even when considering only correct trials, choice representations in the delay were much decreased in the RepL compared to the STM phase, while the Stimulus and Response codes showed robust accuracy increases in both phases (Fig. 5a-c). If trials in the RepL phase were characterized by a transition from cognitive engagement to disengagement as the delay progressed, the Delay code accuracy should be comparable to the STM phase early in the delay, and should then show a drop as delay increased. However, the decrease in delay encoding was uniform throughout the delay period (Fig. 5c,h), indicating that ALM dynamics in the RepL phase were predetermined from the trial’s onset and showed minimal or no loading of memory following stimulus presentation. Consistent with this view, we found individual sessions in which the delay code in correct RepL trials was not significantly above chance at any point in the delay (Fig. 5d). The disappearance of a significant delay encoding during RepL trials was also found in the average activity of single neurons (Fig. 5i) as well as at the population level in single trials (Fig. 5e-f). Moreover, when analyzing errors in the RepL phase, there was no significant trace of code reversals throughout the delay as found in STM error trials (Supplementary Fig. 9a-b). We looked for other neural traces in ALM that could explain the tendency to repeat in RepL trails. The occurrence of attraction biases in delayed-response tasks in human and non-human primates has been related to the encoding of previous memories in the pre-stimulus period (Barbosa et al. 2020). We found that the previous response left indeed a decodable trace that covered the inter-trial-interval but its magnitude at the stimulus onset in the following trial was not significantly different between STM and RepL trials (Fig. 5g). Furthermore, we could not identify alternative neural ensembles to the three codes identified in STM phases which would become active only during RepL trials (Supplementary Fig. 9c-d). Together, these results show that during RepL trials choice encoding during the delay was strongly and uniformly reduced and its temporal dynamics did no longer show signatures of forgetting during error trials, suggesting that choices in these trials were no longer steered by ALM activity.

**Figure 5.**
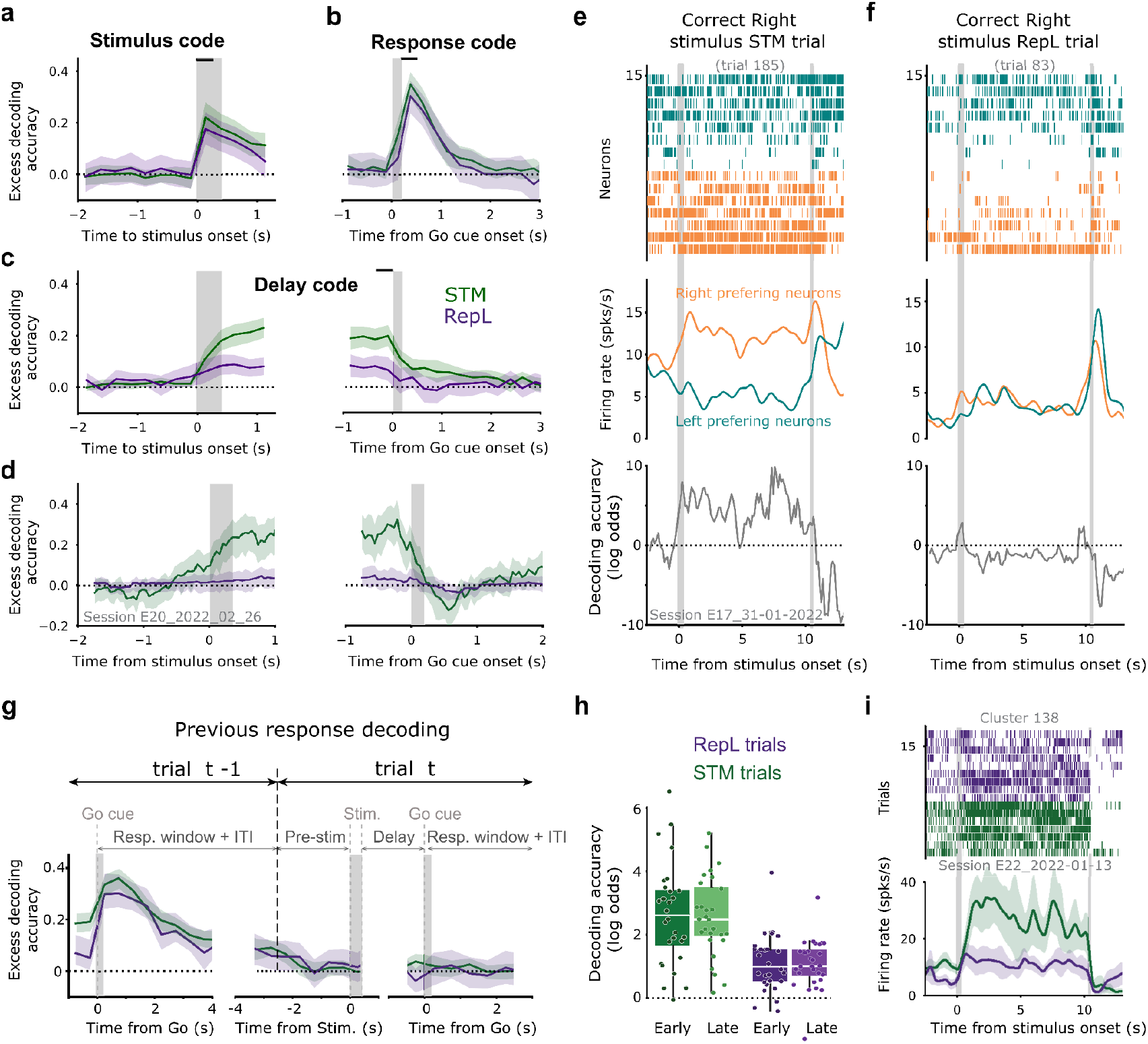
Population encoding during Repeating Lapse phases. **a-c**, Time evolution of the average cross-validated excess decoding accuracy of the Stimulus (a), Response (b) and Delay code (c) tested using correct trials in STM (green) and RepL phases (purple). The Delay code was aligned at both the delay onset (left) and offset (right). Gray shaded areas indicate Stimulus and Go cue durations. Horizontal black bars above indicate the time bin used to train each code. Correct trials with delays 1, 3 and 10 s were used for the training and testing. **d**, same as in (c) for an example session. **e-f**, Examples of two correct Right-stimulus trials, taken from the STM (f) and RepL (g) phases in the same session. Instantaneous activity from Delay code neurons showing selectivity for either Left (blue, n = 7) or Right (orange, n = 7) choices is displayed as in Fig. 4g-h. The RepL trial (g) shows absence of persistent encoding during the entire delay. **g**, Average cross-validated accuracy of previous response from decoders trained on successive time points throughout the trial using all after-correct trials (i.e. previous choice was rewarded). Testing was done on each corresponding time bin, separately for STM and RepL phases. Previous response was decodable from the Go cue of the previous trial (left panel), during the ITI and until the end of the trial when ports moved away at t=-2.5 s (vertical black dashed line). Then decoding accuracy went to zero, and remained zero throughout the delay, for both STM and RepL phases (right panel). **h**, Log odds obtained from the Delay code averaged early (first 0.5 s of delay) vs. late (last 0.5s) tested using both STM and RepL correct trials with delay 1, 3 and 10 s. Dots indicate individual sessions. Linear mixed model: STM/RepL t-value = 28.471, p = 2 · 10^−16^; Early/Late t-value = 2.42, p = 0.06 and interaction t-value = -1.689, p = 0.09. **i**, Spike rasters (top) and average PSTH for correct Left 10s-delay trials occurring in STM and RepL phases from an example neuron.

### Population synchrony increases during RepL states

Lastly, we turned to investigate whether the switching between STM and RepL phases correlated with changes in network synchrony, a metric commonly used to monitor changes in brain state (Jacobs et al. 2020; Einstein et al. 2017; McGinley et al. 2015; Reato et al. 2023). As reported previously (Jacobs et al. 2020; Reato et al. 2023), synchronized spiking activity across the recorded neurons was sporadically but clearly observed in some trials (Fig. 6a). We defined the synchrony index *Synch* for each trial as the normalized standard deviation of the population instantaneous rate (Methods). Thus, Synch was close to one when the network was desynchronized and increased as neurons became more synchronous. To avoid the interference of the stimulus and the response with the circuit’s state, we focused on 2 s of baseline activity before stimulus onset. Within the session, Synch fluctuated broadly being low for most of the session but exhibiting transient surges, reflecting periods of elevated synchrony, particularly (but not only) towards the end of the session (Fig. 6b; Supplementary Fig. 10.). We asked whether these surges in network synchrony were related to RepL phases identified using our behavioral HMM. Indeed, synchrony was significantly larger in RepL phases compared with STM phases (1.29 ± 0.25 and 1.46 ± 0.39 for STM and RepL, paired t-test, t = -3.928, p<0.001, n = 31; Fig. 6b-c). The across-trial correlation between Synch and the model’s posterior probability p(STM) was also significantly negative, even after correcting for the average systematic drift of these variables within each session (corrected correlation coefficient, r = −0.07 ± 0.12, t-test, t = -3.137, p<0.01; Fig. 6d). Consistent with this, the running average of accuracy and RB also correlated significantly with Synch (corrected correlation coefficients, r = −0.09 ± 0.10, t = -4.832, p <0.001 for accuracy; r = 0.05 ± 0.06, t = 4.356, p< 0.001, for RB; n = 31 sessions, Fig. 6d). Finally we wondered whether this increased synchrony was associated with a particular oscillatory frequency band. Autocorrelograms and power spectra of the population instantaneous rate in a 2-s pre-stimulus period separately averaged for STM and RepL trials showed marked differences: during RepL trials, spiking activity showed traces of oscillatory activity which were largely absent in STM trials (Fig. 6e; Supplementary Fig. S13). Averaged ratio of power spectra was significantly above one in the band 3.1 - 15.6 Hz (*t*-test, p<0.05) and peaked at 4.4 Hz (Fig. 6f; median peak frequency was 4.91 Hz with [4.06, 7.88] Hz quartiles, Supplementary Fig. S13; n = 31 sessions; Methods). Thus, the RepL phase tended to coincide with epochs of elevated synchronous, oscillatory spiking in ALM, an activity pattern commonly observed in cortical recordings in head-fixed behaving mice (Jacobs et al. 2020; Reato et al. 2023).

**Figure 6.**
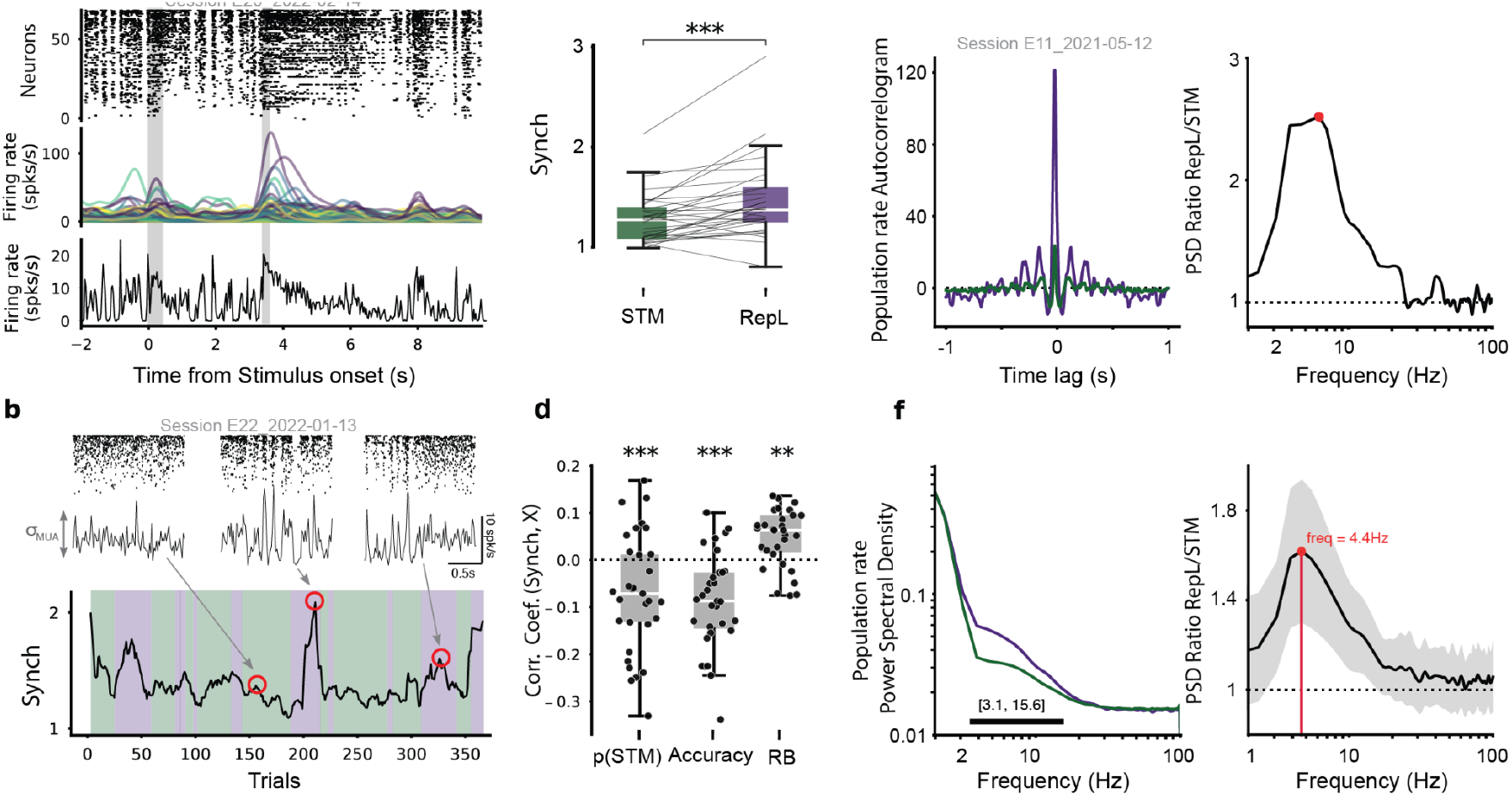
Population synchronous activity occurs more frequently during RepL phases. **a**, Population activity during an example 3s-delay trial with clear synchronization among neurons during pre-stimulus, delay and post-response periods (compare with the trial from the same session shown in Fig. 3a). Top raster shows spike times from 98 simultaneously recorded units, sorted according to their firing rate during the delay. Middle plot shows instantaneous rates for each unit obtained by convolving spike trains with a Gaussian kernel (σ = 150 ms). Bottom panel shows the population instantaneous firing rate, obtained from the merge of all the simultaneously recorded isolated units (20ms bin width). **b**, bottom plot: Synch, defined in each trial as the std. dev. of the population instantaneous rate during the 2 s pre-stimulus period normalized by the std. dev. of the rate obtained if the neurons fired independently (see Methods), for an example session. Background shadings represent STM (green) and RepL phases (purple) identified by the HMM. Top insets show the pre-stimulus population raster and population instantaneous rates of three representative trials (arrows indicate the trial position). **c**, Synch values separately averaged for STM and RepL phases. Each connected pair of dots represents one session (paired t-test, t = -3.93, p = 4.66·10^−4^, n = 31 sessions). **d**, Correlation coefficients between the Synch trace and the posterior p(STM) trace given by the HMM, the binary outcomes (accuracy) and the repeated choices (RB). Systematic within-session temporal drifts were corrected by subtracting session-shuffled correlation coefficients (Methods). Each dot represents one session (t-tests t = -3.14, -4.83, 4.36; *p* = 0.003, 3.74·10^−5^ and 0.0001, resp.; n = 31). **e**, Autocorrelograms (left) and ratio of power spectral densities computed from the population instantaneous rate during the pre-stimulus period in the RepL and STM phases (right), shown for an example session (see Supplementary Fig. S13 for all sessions). **f**, Session-averaged power spectral density separately computed in RepL and STM phases (left) and their ratio (right; n = 31; error band shows 95% C.I.). Horizontal black bar indicates the range where the ratio is significantly different from one (two-tailed, *t-*test, p<0.05).

## DISCUSSION

To characterize the neural mechanisms underlying failures in STM we first dissected mice behavior during a delayed-response task with variable delay. We found that choice accuracy decreased with delay length, a marker of faltering memory maintenance (Fig. 1b-c), but the tendency to repeat previous choices did not (Fig. 1f-g). To reconcile these two features, we used a two-state Hidden Markov model that parsed each session into phases that we interpreted as trial episodes where (1) animals exercised cognitive control so that their responses were guided by the stimulus and depended on STM (control-capable trials), and (2) trials where control was limited and animals just tended to repeat their last action (control-limited trials). We used the model to analyze ALM neural activity in this task and found that, during engagement phases, ALM population dynamics had all the features of an attractor network both in correct and error trials, whereas during task-disengaged phases, the same populations exhibited very different dynamics with a lack of clear sustained memory activity and enhanced synchronous activity in the circuit.

We aimed to identify the causes of STM forgetting errors, defined as those caused by the disruption of memory during the maintenance period. The occurrence of this type of errors was revealed by the decrease of the accuracy as a function of the delay duration (Fig. 1c). By separating the trials into control-capable and control-limited phases we were able to identify RepL trials, where errors did not seem to be caused by memory maintenance, and focus on STM phases. We found that, in error STM trials, the population of neurons maintaining the encoding of the prospective response during the delay started encoding the correct choice at the delay onset but ended up encoding the wrong one after delays of D = 10 s (Fig. 4c-e). These code reversals at long delay intervals demonstrate that, at least in a significant fraction of error trials, the circuit fails to maintain the correct memory initially kept in STM and transitions into the opposite, wrong memory representation. Whether reversals reveal punctuated rapid transitions occurring at moments randomly distributed along the delay (as in our model, Supplementary Fig. 7e-g), or a more slow degradation spanning hundreds of milliseconds remains an open question, as both are compatible with the average reversal trace that we found (Fig. 4c) (Latimer et al. 2015).

Previous work using a similar task has demonstrated DW attractor dynamics during the delay period in ALM (Inagaki et al. 2019; Finkelstein et al. 2021a; Yang et al. 2022). Crucially, when ALM population activity was perturbed by transient photo-inhibition in the early delay, the activity quickly recovered to encode the upcoming response as in unperturbed trials (Inagaki et al 2019). This implies that, once released from the photoinhibition, the circuit evolved rapidly towards one of the two only seemingly possible states associated with the two responses. Our results are consistent with this view of ALM operating as a bistable attractor network and extend it by showing that the network dynamics can generate spontaneous attractor transitions during long delays. However, none of these previous studies found evidence supporting this fundamental prediction of attractor dynamics when they looked at unperturbed spontaneous error trials. Instead, neurons along the delay code axis seemed to encode the wrong response even before stimulus presentation (Yang et al. 2022; Inagaki et al. 2019). The fact that reversals in our data were revealed only at the longest delay D = 10 s (they used D = 0 - 4 s) and that we had to remove RepL error trials to isolate reversal-type errors can probably explain the difference between studies. Taken together, our study supports the existence of two attractors along the dimension of the Delay code and suggests that stochastic-driven transitions between these attractors is a common mechanism underlying STM forgetting.

Attractor network models describing sustained short-term memory activity generally operate in what is called the multi-stability regime in which, in addition to the attractor states associated with the remembered items, there is always a stable baseline state with low homogeneous spontaneous firing across the network (Amit and Brunel 1997; X. J. Wang 1999; Hansel and Mato 2013; Brunel and Wang 2001; Compte et al. 2000). In such a regime, the exact same network (i.e. without modifications in its inputs or connectivity) can reproduce the sustained activity observed during mnemonic periods as well as spontaneous activity characteristic of periods without memory maintenance. Input fluctuations during the mnemonic period can then impact memory encoding in different ways, causing transitions between memory states (i.e. memory reversals) or between memory states and the spontaneous baseline state (i.e. memory loss). In multi-stable discrete attractor networks, transitions between attractors are typically dominated by transitions between memory states and the baseline state, as the latter is typically centrally interposed in state space (Renart et al. 2007; Wong and Wang 2006). Hence, the prediction from a discrete attractor model in the multi-stability regime is that memory forgetting would generally take the form of memory loss rather than memory reversals. Memory loss would yield average decoding traces slowly degrading towards zero rather than the reversing pattern we observed (Fig. 4c), which suggests that the ALM circuit is not operating in this multistable regime in our data. With slight changes in parameter values, these same network models can operate in the winner-take-all regime (WTA) where there is no longer a spontaneous activity stable solution and the system’s only fixed points are the memory attractors (Roxin and Ledberg 2008). The DW model exemplifies this WTA regime for the case of two attractors (Prat-Ortega et al. 2021) and can cause memory reversals but not memory loss. Importantly, using the WTA regime for STM requires that the inputs or connectivity of the network change from the prestimulus period to the delay period to maintain persistent activity. Previous work has proposed that an unspecific (i.e. carrying no information about the memory being held), excitatory external drive starting at stimulus onset and spanning the delay period can make the circuit undergo this transition (Inagaki et al. 2019; Itskov, Hansel, and Tsodyks 2011; Finkelstein et al. 2021b). Furthermore, when delay duration is fixed and the response timing can be predicted, the external drive ramps up giving rise to ramping delay activity, whereas when delays are variable, the external drive is constant giving rise to sustained stable activity (Inagaki et al. 2019). We propose that this external drive to ALM reflects the level of cognitive control, so that it engages or disengages this frontal area from STM function. The necessity of such a control-dependent signal for attractor maintenance opens the possibility that behavioral errors are caused by the *exhaustion* of this input as the delay progresses. The signature of this forgetting-by-exhaustion mechanism would be a decrease towards zero of the delay activity encoding the correct choice and not the reversal from correct to error that we observe. Our data however does suggest that changes in cognitive control across the session could correlate with changes in the amplitude of the external drive, as we discuss below.

Recently, there has been an increased interest in characterizing changes in behavioral state across trials during decision making tasks, particularly in mice (Ashwood et al. 2022; Bolkan et al. 2022). One study found that mice performing an accumulation of evidence task in a virtual T-maze alternated between three states: a state where decisions were driven by the accumulated stimulus evidence, a state in which the stimulus had almost no impact but previous responses did, and a third state intermediate between these two (Bolkan et al. 2022). Our behavioral analyses are consistent with this previous work, where we have in addition included a more mechanistic description of the task-engaged state and we have identified the task-disengaged state as history-dependent lapses. However, the crucial advancement of our study is to shed light on the neural correlates of the behavioral strategies in two important ways. First, our decoding analysis provides evidence that the dynamics of ALM in representing and maintaining the upcoming response during the mnemonic period are fundamentally different between the two states (Fig. 5). Second, we found that phases of neural synchronization in the ALM tended to coincide with phases of task disengagement (Fig. 6).

During awake behavior, subjects transition between brain states typically characterized by different degrees of synchronous activity measured as large, low frequency fluctuations in either the local field potential, the intracellular membrane potential or the population spiking activity (Einstein et al. 2017; Jacobs et al. 2020; Beaman, Eagleman, and Dragoi 2017; Engel et al. 2016; Reimer et al. 2014; McGinley, David, and McCormick 2015; Sachidhanandam et al. 2013; Crochet and Petersen 2006). Whereas factors such as running, whisking and pupil dilatation have been shown to systematically covary with brain state (Reimer et al. 2014; McGinley, David, and McCormick 2015; Vinck et al. 2015), the relationship between brain state and behavioral performance in decision making tasks is still debated. Some studies have found that cortical synchronization had no significant impact in a detection (Sachidhanandam et al. 2013) or a Go/NoGo task (Einstein et al. 2017). Another study carrying out a visual two-alternative choice task, found a specific impact on engagement (higher synchrony causing more misses) but not on perceptual discriminability (Jacobs et al 2020). More recently, Reato and colleagues have found that elevated cortical synchrony can reduce discrimination performance in a two-alternative forced-choice auditory task but only if the previous outcome was an error (Reato et al 2023). The authors explain the absence of impact of synchronous activity by arguing that “when discriminating simple sensory stimuli, mice can operate in a behavioral state equivalent to ‘flow’ during streaks of correct trials”. Supposedly, the cortex in this state may not play a major role in controlling these more automatic responses. An error however, represents an unexpected signal which recruits performance monitoring by areas such as the anterior cingulate cortex (ACC) (Norman et al. 2021). In contrast with this study and the above-mentioned studies, our results did show a correlation between trial-to-trial synchrony and choice accuracy, without having to condition on previous outcome (Fig. 6d). A possible explanation for this difference may be the higher cognitive load of our task in which mice had to maintain their responses during long and variable delays, a condition that may require a more sustained control by frontal circuits. This is consistent with results from another STM study, in which small increases in population synchrony in the visual cortex during the delay tended to cause a decrease in animals’ performance (Beaman, Eagleman, and Dragoi 2017). While in this study the decrease in performance was associated with the worsening of the encoding of the target stimulus in the visual cortex, we found that high synchrony in ALM was associated with RepL phases (Fig. 6c) during which persistent activity was severely reduced (Fig. 5c-d, h). This does not mean however that population synchrony had a direct impact on the stability of persistent states, as we saw no traces of memory maintenance errors in RepL trials (Supplementary Fig. 9a). Instead, our data suggest that synchrony may be reflecting changes in cognitive control that alter the dynamics of ALM and its role in guiding behavior as discussed above.

Following this reasoning, switches between STM and RepL phases could reflect the need to periodically relieve the effort that sustained control of cognitive behavior requires (Botvinick et al. 2001; Shenhav et al. 2017). It is well known that tasks that rely on some form of STM commonly require high levels of cognitive control and that the frontal cortex plays a major role in exerting it, such that in its absence responses can be impulsive or automatized. In our task, the water ports moving close to the mouse’s snout act as a very strong cue for the animal to respond. Previous work has characterized an excitatory input from thalamus to ALM as carrying this Go signal which ultimately triggers the activation of the response neurons (Inagaki et al. 2022). Our results suggest that this Go signal into the ALM is present both in STM and RepL trials, as the animals respond to the approaching of the ports during the two phases. The difference between control-capable and control-limited trials would be the state of the network along the Delay code axis set by the unspecific external drive mentioned above: in STM trials, ALM delay neurons display sustained activity that causes the response to the Go signal to be steered towards the side showing persistent activity (Supplementary Fig. 11a). In contrast, in RepL trials, this sustained activity is absent, and the Delay code neurons do not exert any control over the animal’s response to the Go signal which would depend on a different tiebreaker mechanism (Supplementary Fig. 11b). In this interpretation, repeating lapses reflect the limits of this control and should be construed as the default control-free policy (Castiñeiras and Renart 2022), in contrast to other proposals such as reflecting exploration (Pisupati et al. 2021) or the misbelief of animals about the environment’s stationarity (Gupta et al. 2024). Our interpretation moreover agrees with the observation that towards the end of the session animals exert less control (Supplementary Fig. 2b), possibly because of a devaluation of the water reward as mice get satiated or an increased sensitivity to effort as they get mentally tired (Shenhav et al. 2017). Because lapses were strongly influenced by previous responses (Fig. 2i-j), but were independent of previous outcomes (Supplementary Fig. 3d-e), the repeating bias may reflect a process of value-free *action reinforcement (Akaishi et al. 2014; K. J. Miller, Shenhav, and Ludvig 2019; Greenstreet et al. 2022)* (Supplementary Fig. 11b). Following this rationale, animals would learn the relevant stimulus-reward associations of the task via some instantiation of value-based reinforcement learning, whereas they would consolidate the learned behavior by the reinforcement of recent stimulus-action associations independently of reward outcome (Greenstreet et al. 2022; K. J. Miller, Shenhav, and Ludvig 2019). In that respect, the repeating tendency of the lapses in our task would be the residual signature of the underlying reinforcement of the association between Go cue and Right or Left licking, a process possibly occurring across all trials but only detectable during phases of cognitive control withdrawal (Supplementary Fig. 11b). Future work should aim to characterize the circuitry underlying the repeating bias and its interaction with cognitive control. Our results indicate that cognitive control mediates the intermittent recruitment of attractor dynamics in ALM allowing the maintenance of STM which ultimately serves to decouple our decisions from both the immediacy of the sensory world and the inertia of our previous actions.

## Acknowledgments

We thank Carles Sindreu, Klaus Wimmer and Lorenzo Fontolan for valuable comments on the manuscript. We thank Balma Serrano, Eva Carrillo and Aina Fenández Rodríguez for animal training. Also, thanks to the members of the Brain Circuits and Behavior lab (IDIBAPS) for fruitful discussions. This research was funded by Fundación privada Cellex, Instituto Carlos III (PIE 16/00014), Spanish Ministry of Economy and Competitiveness with the European Regional Development Fund (SAF2015-70324-R), Fundación La Caixa para Doctorados en España 17/18 (L-CF/BQ/DE17/11600009), by MCIN/AEI/ 10.13039/501100011033 and by “ERDF A way of making Europe” (Grant PID2021-126698OB-I00), by CERCA Programme/Generalitat de Catalunya, and by Generalitat de Catalunya (AGAUR 2014SGR1265, 2017SGR01565). This work was developed at the buildings Centre Esther Koplowitz and Centre de Recerca Biomèdica Cellex (IDIBAPS, Barcelona). J.R. appreciates the hospitality of the Grossman Center for Quantitative Biology and Human Behavior at the University of Chicago.

## Author contributions

T.O.-J., A.C. and J.R. designed the experiments with the assistance of C.L.; T.O.-J. carried out all the experiments, analyzed the data and made the figures. G.P.-O. developed the model. T.O.-J., A.C., and J.R. interpreted the data. J.R. wrote the first draft of the manuscript. T.O.-J., G.P.-O. and A.C. contributed in the writing of subsequent versions of the manuscript. All authors revised the final manuscript and gave critical comments. A.C., J.R. and J.D. obtained the funding.

## METHODS

### Animal subjects

Wild type C57BL/6J male mice (n = 48, 18-26g) with initial ages between 8 to 12 weeks were used in this study. They were kept in stable conditions of 23º and humidity of 60% with a standard 12 h dark-light cycle. Mice were housed in groups of up to 4 animals unless surgical or implant conditions required otherwise, in which case they were individually housed. Animals’ cages were environmentally enriched with one running wheel and one plastic tube. Mice had unlimited access to food but they were water restricted for the entirety of training and recording. Prior to water restriction, baseline weight was collected during three days and averaged to give target animal weight. At all times, mouse weight was kept higher than 80% of that initial value by providing water supplements and special care or even by removing the animal from water restriction. Mice always obtained a daily amount of 30 μl of water per gram, which they usually obtained during the experimental sessions. During the weekends or non-working days, mice were given ad-libitum access to water with a 4% concentration of citric acid, or they were manually fed with their required doses of water for the day. All experiments included in this work were conducted according to the Comité d’Experimentació Animal of the University of Barcelona (33/20).

## Behavioral Task

We designed an auditory two-alternative delayed choice (2ADC) task (Erlich, Bialek, and Brody 2011); Guo et al. 2014; Li et al. 2015) with varying delay duration (Fig. 1a). Mice were head-fixed with two speakers located on each side of their head at 2 cm from each ear. Each trial started with the retraction of the water spouts, which were kept out of reach from the mouse until the response window (Supplementary Fig. 12). After port retraction, there was a fixed pre-stimulus period of 2.5 s (3 s in some animals). After this period, a cue was presented consisting of broadband noise (2-20 kHz) at 72 dB played during 400 ms from one of the two speakers, Left or Right. After the sound cue, a delay was imposed, randomly selected from a set of discrete values *D* = 0, 1, 3, 10 s (Groups #2 and #3). One group of animals were trained with only *D* = 0, 1 and 3 s (Group #1; see Table 1). Following this delay, a motor approached two lickspouts, cueing the mouse to respond by licking one of them. If mice licked the port corresponding to the side of the sound cue, they obtained a water reward of 2.5 μl. If they licked the opposite port, no reward was provided and a weak LED light turned on during a 2 s timeout (Supplementary Fig. 12b). If no response was given during the response window (2 s), the trial was considered invalid, neither reward nor timeout were provided and it was excluded from the analysis (average 4% ± 2%). After any of these outcomes (i.e. correct first lick or end of the timeout), an intertrial interval (ITI) of 4 s was introduced until the beginning of the following trial (i.e. port retraction). To facilitate mouse task engagement at the session start, we increased the fraction of easy trials with zero delay during a “warm-up” period (fraction was 0.8 in trials 1-20 and 0.5 in trials 20-40; in the rest of the trials it was 0.25, like the rest of the delay values). Warm-up trials were excluded from the behavioral analyses shown in Figs. 1-2. Left and Right stimuli were presented randomly with equal probability, without any serial correlations across trials. Sessions ended after 60 minutes or when animals did 15 invalid responses in a row, whatever happened first.

### Behavioral training

Behavioral training involved four stages: Habituation, Lick teaching, Categorization learning and Delayed-response learning. During the first 3-5 days and before water restriction, animals were habituated to the experimenter and to head fixation. After this, they were placed under water restriction and Lick teaching began. This step involved animals collecting water freely from the lickspouts inside the behavioral box while remaining under head fixation. Mice were rewarded every time they spontaneously licked either port. No other cues such as lickspout movement or sounds were present in this step. Once they could collect reward from both lick-spouts (1-3 days), they progressed to the trial-structured Categorization learning stage, where they were trained to associate the sound location with the rewarded side. In each trial, a lateralized sound was played for 2 s or until the mice licked the correct port. To help animals create the association, during the first days (1-3 days) they were allowed to correct error responses as the protocol ignored incorrect licks. After that, animals were punished for licking the wrong port with a 2 s timeout. In addition, a fixation window with no licking at trial onset of up to 2 s was required before the stimulus could be played to discourage impulsivity. Finally, in these stages, a correction bias was implemented to correct the strong preference of some mice to lick mostly one port, consisting of an increased probability for trials on the non-preferring side during epochs of high bias (60-40% maximum asymmetry). When animals achieved an accuracy above 90% during 2 consecutive days (5 to 7 days) they advanced to Delayed-response learning. Training in this stage aimed at teaching animals to delay their responses after the stimulus presentation. We used a motor to retract the lickspouts away from the animals’ reach, to teach the animals when to lick and when not to lick. First, animals were habituated to the motor by retracting the ports after the ITI, for 2.5 s (pre-stimulus period) before stimulus onset, and returning them right after stimulus offset (5-7 days). Then, delays were introduced by returning the ports later and later after stimulus offset (Fig. 1a). Trials in each session were always classified in three different delay lengths, presented randomly. Initially, these delay lengths were set to zero and were progressively increased within and across sessions if the animal performed well until reaching the values 0, 1 and 3 seconds (7 days). Delays of 10s were added after animals achieved robust accuracies with delays 0, 1 and 3s for more than two weeks. At the end of the training, all delays were presented with the same probability, except during the warm-up trials at the beginning of each session (see above). Delay 0 seconds actually lasted 100 milliseconds, a period imposed to guarantee that stimulus presentation did not overlap with motor movement.

### Behavioral apparatus

The behavioral task was performed inside a custom-made acoustic isolation box built with high-density insulation foam (Copopren; isolation around 40 dB). The majority of the components used were either 3D printed using ABS-like resin or PLA or anodized steel parts (Thorlabs). The main behavioral state-machine was controlled using Bpod (Sanworks), with two extra Arduino boards (UNO and Mega) controlling sound delivery and the stepper servo motor. Acoustic stimuli were delivered through two mobile-phone speakers (ZT-026 YuXi, Aliexpress). Sound stimuli were played using an Arduino and an Adafruit “MP3” Music Maker Shield (codec VS1053B), playing sounds previously loaded on an SD memory card with a frequency rate of 48 kHz. The water delivery and lick detection were provided by a custom-made lickspout with two tubes separated by 6 mm, making an angle of 90º and 18 gauge. Licks were detected using two infrared photogates with beams finely calibrated to transverse the licking path to each water tube. Water-port position was adjusted with a 5V linear stepper servo motor. The water reward was delivered using two 12V peristaltic pumps that were calibrated weekly. One visible-light LED was placed in front of the animal to cue error responses. Animal head-fixation was performed outside of the boxes, using a portable caddy (Thorlabs) with custom-printed steel holders (Guo et al. 2014). All the sessions were recorded using infrared cameras (ELP 1080P CMOS OV2710).

### Surgeries

Animals underwent headplate implantation surgery before training started. Mice were deeply anesthetized in an induction chamber with 2% continuous isoflurane flow and 0.8L/min of oxygen concentration. When the animal was unresponsive to paw pinch and showed a regular breathing pattern, it was transferred into a stereotaxic frame (Kopf Instruments) and isoflurane levels were kept between 1-2% for the rest of the surgery. Antibiotic ointment (Cusi Antiedema, 50mg/g) was placed and maintained on the animal’s eyes during the procedure. Body temperature was regulated using a rectal temperature probe and a homeothermic blanket (Stoelting). Buprenorphine (0.1 mg/kg) was applied subcutaneously. The fur on the head was cut using scissors and the scalp cleaned and disinfected using Ethanol (70%) and iodine antiseptics (Betadine) with cotton swabs. After cutting and removing the skin, Lidocaine (0.8%) was applied and the connective tissue and blood was carefully removed using peroxide acid (H_2_O_2_). A scalpel blade was used to further scratch the skull surface and remove any attached tissue. Lambda and Bregma were z-aligned, four anchoring screws (ComponentSupply, USA) were inserted and the steel headplate (adapted from Guo et al. 2014) was glued between Lambda and Bregma using Vetbond glue. Once dried, dental cement (Paladur) was applied covering all the exposed area and the top of the implant. Once the cement was dry, mice were injected with saline (1 ml subcutaneously) and transferred to an individualized recovery area with wet pellets and a heat blanket where they were carefully supervised until recovery from anesthesia. During the following days, their weight was monitored and daily doses of analgesia (Meloxicam, 2mg/kg) were administered. Water bottles included an antibiotic (Enrofloxacin, 100mg/L) which was removed 2 days after surgery if no signs of disease were observed. Training started at least one week after headplate implantation.

Apart from the standard surgical procedure for headbar implantation, mice undergoing electrophysiology experiments had two further modifications. First, ALM coordinates +2.5AV ± 1.5M (Guo et al. 2014) were marked and a really thin layer of transparent cement applied on top of it. Additional buildup of cement was placed surrounding the coordinates to create a well-like area. For grounding, two separate screws soldered to an electronic pin were placed on top of the cerebellum. The pin providing the best ground was later used for recordings.

### Electrophysiological recordings

Electrophysiological recordings were performed in mice from Group 4 (n = 8) using silicon probes either from NeuroNexus (Buzsaki64 rev4 or A4×16-Poly2-5mm-20s-150-160) or Neurotech (ASSY-77 H6, 2 shanks). Probes were coupled to an adapter (Neurotech Adpt-A64-OM32×2) and then attached to a micrometric oil manipulator (Kopf Instruments). The collected signal was amplified and digitized using preamplifier boards (RHD2132, Intan Technologies). The signal was collected using an Open-ephys acquisition board (Siegle et al. 2017). This same board also received two TTL signals (delay onset and stimulus identity) through a breakout box (LabMaker, multichannel I/O board) coming from the Bpod and further used for alignment and trial identification.

The evening before the beginning of the recordings, animals were briefly anesthetized and fixed in the stereotaxic frame. Using a dental drill, a craniotomy (2 mm diameter) and durotomy in one hemisphere was performed centered around the marks that were drawn during the headbar implantation surgery. The area was carefully cleaned using cotton swabs and sugi sponges, removing any bone or cement debris. When finished, a layer of Kwik-Cast (World Precision Instruments) was used to seal the craniotomy.

On the day of the recording, the Kwik-cast plug was removed, the craniotomy was rapidly cleaned and a probe was lowered 500μm to 1200μm below the cortex surface (ca. speed 1-5μm/s). After placement of the electrode, a drop of agarose with aCSF (1%) was added embedding both the craniotomy and the shanks. The brain was allowed to settle from 10 to 20 minutes before starting the behavioral task and the ground pins were connected to the probe. 3 to 5 penetrations per craniotomy were performed on average (range 6 - 10 per animal).

Recordings were visualized online using Open Ephys software. Broadband activity was sampled at 30kHz per channel, digitized and band-pass filtered between 2 and 6500 Hz and then stored for offline analysis. Raw data was then spike-sorted using Kilosort version 2.0 (github.com/MouseLand/Kilosort2). Units were manually curated using Phy (github.com/cortex-lab/phy). Visual inspection of contamination indexes, auto and cross correlograms of clusters, PCA projections and overall shape of spike waveforms was performed and following such clusters were either discarded or classified as multi (MUA) or single unit activity (SU).

### Behavioral analysis

All measurements reported in the text refer to mean ± SD unless indicated otherwise.

### Repeating Bias

We defined the Repeating Bias (RB) to quantify the tendency of mice to repeat the previous choices:

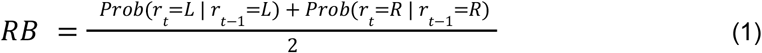

where *Prob*(*r*_*t*_ = *X* | *r*_*t*− 1_ = *X*) is the conditioned probability of making a particular response *r*_*t*_ = *X* when the previous choice r_t-1_ had also been *X* (with *X =* Left or Right). The RB can take values above 0.5 indicating that there is a tendency to repeat the previous choice or below 0.5 indicating there is a tendency to alternate. Computed this way, the RB controls for fixed choice biases independent of previous responses since in that case the conditioned equals the unconditioned probability, i.e. *Prob*(*r*_*t*_ = *X* | *r*_*t*− 1_ = *X*) = *Prob*(*r*_*t*_ = *X*), giving rise to *RB* = 0. 5.

### Autocorrelation analysis

Autocorrelograms were used for characterizing the temporal correlations in the binary sequences of (1) choice outcomes and (2) repeated responses (Weilnhammer et al. 2023). We first obtained the raw autocorrelograms in each session and averaged it across sessions for the same mouse. To control for the slow trend in accuracy and repeating bias observed across the session (Supp. 1), we calculated the cross-correlogram between outcome or repeat sequences taken from two different sessions, averaged across many pairings (from 20 to 80 surrogate sessions, always corresponding to the same mouse). The corrected autocorrelogram was finally obtained by subtracting the averaged cross-correlogram from the averaged autocorrelogram. This correction allowed us to quantify short-lived dependencies across trials that were not simply caused by the slow global non-stationarity of these observables.

### Generalized linear mixed models

We fitted a GLMM to quantify the influence that the different task variables had on the choices of each individual mouse (Busse et al. 2011; Frund, Wichmann, and Macke 2014; Abrahamyan et al. 2016; Braun, Urai, and Donner 2018; Hermoso-Mendizabal et al. 2020; Reato et al 2023). We used both mouse Groups #1 and #2 together in this analysis. The binomial response variable *r*_*t*_ (*r*_*t*_ =1 for Right choices and *r*_*t*_ =-1 for Left choices in trial *t*) was fitted using a logit link function as a combination of the following task regressors (see Fig. 1k and Supplementary Fig. 2e):

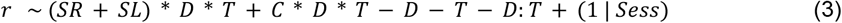

The current stimulus was represented by the binary regressors *SL* and *SR* that indicated whether the current stimulus was Right (*SR* = 1 and *SL* = 0) or Left (*SR* = 0 and *SL* = 1). The delay regressor *D* took the values 0, 1, 3 or 10 of current delay. The Action Trace *C* is defined as a weighted sum of previous choices, independently of their outcome, which tracks which is the most abundant choice in the recent trial history:

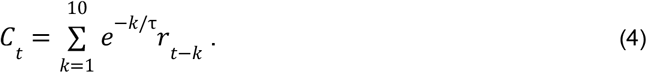

The decay constant of the Action Trace was obtained independently for each mouse from fitting exponentials to the decay of the weights of previous choices (average τ = 2. 1 trials, see Supplementary Fig. 1). The trial index regressor *T* transformed the trial index in each session to a continuous variable in the interval [0,1] with 0 corresponding to the first trial and 1 to the last. The weights associated with the *T* regressor are shown in Supplementary Fig. 2f-i. Finally, the grouping variable session *Sess* was taken as a random intercept that controlled for day to day variations in preferred choice side. According to equation (2), the model had 11 weights for SL, SR, SL×D, SR×D, SL×T, SR×T, SL×D×T, SR×D×T, C, C×D and C×D×T. The four regressors {SR, SL, SR×D, SL×D} can be reparameterized as {S, D, S×D, 1} with *S= SR* − *SL=* {+1,−1} for Right and Left stimuli, respectively. Thus, in the chosen parameterization, the regressors SL and SR were collinear, i.e. SL+SR=1, so the model does not include the fixed intercept explicitly. The chosen parametrization was preferred because the idiosyncratic Left-Right biases, often observed in individual mice (Fig. 2q-r), were easily interpretable from unbalanced weights of the regressors SL and SR, and all their interactions. For example, the weight of SL×D would be related to the transition probability from L→R.

### Repeat-independent double well attractor model

We used a one dimensional diffusion process that, in the winner-take-all regime, represents a mathematical reduction of a classical spiking neural network commonly used to model cognitive processes like short-term memory and decision making (Roxin and Ledberg 2008; Wong and Wang 2006; X.-J. Wang 2002). Thus, for certain parameter values, this one-dimensional system can be viewed as the motion of a noise-driven, overdamped particle in a double-well potential. The dynamics of this particle, whose position we define as the choice variable *x(t)*, is defined by

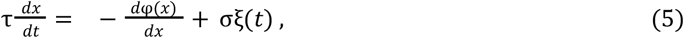

where τ is the time constant of the system, ξ(*t*) is a Gaussian white noise with zero mean and unit variance and the potential is given by

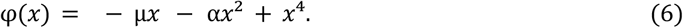

The coefficient of the quartic term is set to one, without loss of generality. Moreover, to simplify the model, we set the magnitude of the white noise to σ = 1 (although the expressions below maintain the explicit dependence for generality), and use the free potential parameters *α* and µ to fit the model to each animal’s behavior. The parameter *α* controls the height of the barrier between the two attractors while µ controls the asymmetry between them. We use the same dynamics defined by Eqs. 5-6 for modeling the two sequential steps of the task, namely (1) memory loading and (2) memory maintenance.

The loading step is modeled as the fast evolution of the system from a fitted initial condition *x*_0_, somewhere near the maximum of the potential, until it reaches one of the two wells (Prat-Ortega et al. 2021). To bias this initial loading state towards the right attractor, the stimulus induces a tilt in the potential towards the corresponding side (Eq. 5) of symmetric magnitude µ = µ _*S*_, where µ _*S*_ = + *S* for the Right stimulus and µ _*S*_ = − *S* for the Left stimulus, and *S* is the fitted parameter. Assuming that the time to initially reach one of the two attractors is much shorter than the time spent on them, we can compute the probability *P*_*R*_ (µ _*S*_) that the system first visits the Right memory upon presentation of the stimulus µ _*S*_ which can be expressed as:

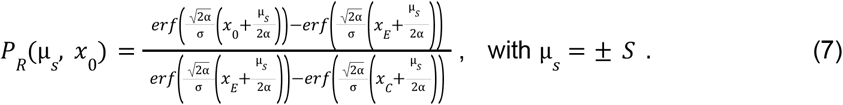

The coordinates *x* _*C*_ and *x* _*E*_ represent the position of the two minima corresponding to the **α** correct and error attractors, respectively, which depend in turn on the potential parameters (, µ) (i.e. *x* _*C*_ is the rightwards minimum of the DW when the stimulus is µ _*S*_ = + *S* and vice versa). By fitting an initial position *x* _*0*_ ≠ 0, the memory loading step can be biased towards one of the attractors above and beyond the bias introduced by the stimulus. Thus, once *S* and *x* are fitted, we summarized their impact in two independent parameters that we defined as the probability to load the Right memory when the Right stimulus is presented *P* _*R*_ = *P*_*R*_ (µ _*S*_ = *S, x*_*0*_) and the probability to load the Left memory when the Left stimulus is presented *P*_*L*_ = 1 − *P*_*R*_ (µ _*S*_ =− *S, x*_*0*_). Notice that, even though our stimuli are symmetric (µ _*S*_ = ± *S*), because in general *x*_*0*_ ≠ 0, these two probabilities are different *P*_*R*_ ≠ *P*_*L*_.

For the memory maintenance during the delay, the Left→Right transition rate *k*_*L*_ and Right→Left transition rate *k*_*R*_ produced by the diffusion noise and can be computed using Kramers rate theory (Kramers 1940):

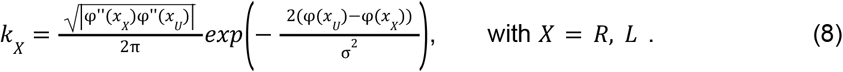

where *x*_*U*_ is the position at the maximum between the two wells (i.e. the unstable attractor state). Note that during the delay there is no asymmetry created by the stimulus (*S* = 0). However, during the delay, the potential can be biased towards one of the possible choices independently of the stimulus and of previous responses, making the two transition rates different. This implies that during maintenance, the asymmetry parameter is in general µ ≠ 0 (Fig. 2a). Using these transition rates, the system can be treated as a Continuous Markov Chain with two states: Left and Right. The probability that at the end of the delay *D* the decision variable is in state Y given that it started the delay in state X is (Durrett 2016):

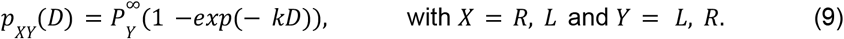

where *k* = *k*_*L*_ + *k*_*R*_, 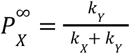 is the stationary probability of finding the system in state *X* when *D* → ∞. Ultimately, our model provides the probability of the system to be found in one choice attractor versus the other at the end of the delay given a Left or Right stimulus. This is achieved by combining the results of memory loading and maintenance described above (Eqs. 7-9) to obtain the probability of being in the *X* attractor in a trial with stimulus µ _*S*_ = ± *S* and delay D, given the model and stimulus parameters:

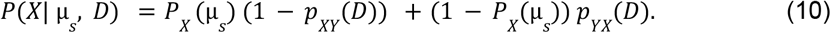

Finally, the choice action after the delay is assumed to be a noiseless process that simply reflects a read out of the memory at that point in time (Prat-Ortega et al. 2021).

### Extended double well attractor models

To investigate whether a double well attractor model (DW) could also exhibit the repetition bias and its dependence with the delay length, we extended the repeat-independent double well attractor model by adding a dependence of the bias parameters *x*_*0*_ and µ on the trial-by-trial value of the action trace *C* (Eq. 4):

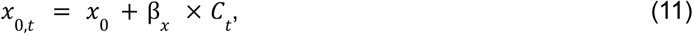

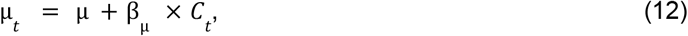

Thus, *C* _*t*_ could bias the memory loading probabilities *P* _*L*_ and *P* _*R*_ (Eq. 11) and/or the transition rates during the maintenance period (Eq. 12). We consider 4 versions of the double well model: (1) the DW without any dependence on *C* _*t*_ (DW); (2) with only the dependence of the loading probabilities *P*_*L*_ and *P*_*R*_ (Eq. 11; *DW-L*), (2) with only the dependence of the asymmetry maintenance parameter µ (Eq. 12; *DW-M*), or (3) with the dependence of both parameters (Eqs. 11-12; *DW-B*; Fig. 2a).

We separately fitted each version of the DW by numerically maximizing the log-likelihood of the data given the model:

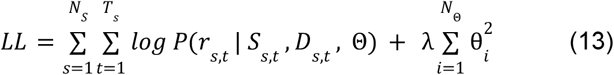

where the function P is given by Eq. 10, with *r*_*s,t*_, *S*_*s,t*_ and *D*_*s,t*_ being, respectively, the animal’s choice, the stimulus and the delay in the *t*-th trial of the *s*-th session, and Θ are the set of model parameters. Here, P is the probability that the model correctly predicted the animal choice given the set of parameters Θ. The second term is the L2 regularization where we set λ = 0. 1 after a grid search where we tried λ = 0. 01, 0. 1, 0. 2, 0. 5, 1. We used gradient descent to find the parameters Θ that maximized the *LL* function for each animal. For this purpose, we used the LBFGS algorithm implemented in the Optim package together with JuliaDiff to automatically compute the gradients. Because in general LL is a non-convex function with respect to the parameters of the DW, the parameters obtained in one gradient descent optimization can correspond to a local maximum. In order to find the global maximum, we ran the algorithm with 100 random initial conditions and we selected the parameters that best fit the data (typically around 2-5% of the initial conditions fell in the vicinity of this optimal solution). To perform model comparison of the various DW models and the Hidden Markov Model (below; Fig. 2e), we used cross-validated likelihood: the data was divided into two halves, the first one used to fit the model and the second one used to compute the likelihood (Eq. 13). Then exchanged the two folds and the likelihood of the other half was computed. The likelihood of the two halves was finally added and divided by the total number of trials and by log(2), to provide the units bits per trial (Fig. 2e). The rest of the panels showing the results from the fitted DW models were not cross-validated.

### Hidden Markov Model

We used a Hidden Markov Model (HMM) with two latent behavioral states, named *Short-term memory* (STM) and *Repeating Lapses* (RepL), such that the subject, in each trial, is in one of the two behavioral states and responds on that trial based on the choice strategy defined by that state (Ashwood et al. 2022; Bolkan et al. 2022). However, the latent variable *z*_*t*_ defining which state is active in each trial *t* is unobservable, a hidden variable, and only the subject responses are observed. The HMM assumes that the state in the next trial depends only on the current state but not on past states. Thus, the parameters of the HMM are the transition probabilities P(STM→STM) and P(RepL→RepL) (Fig. 2d) and the initial probability in trial one to be in the STM state Π = *p*(*z*_1_ = *STM*) (the initial probability to be in the RepL state is *p*(*z* _1_ = *RepL*) = 1 − Π). In addition, each of the two states has its own parameters, which define the responses chosen in each trial.

### RepL state

This behavioral state represents the *control-limited* state in which mice had a limited level of cognitive control and responded to the Go cue by licking one of the two ports but they ignored the stimulus that had been presented in that trial. The probability to make a Right choice in this state is therefore defined independently of the stimulus and only determined by the history of previous choices (Fig. 2d):

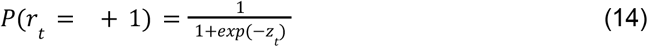

where *z* _*t*_is given by *z*_*t*_ = β _*C*_ *C*_*t*_ + β _*0*_, β _*C*_ is the weight of the Action Trace *C*_t_ defined in Eq. 3 and β _*0*_ is an overall choice bias.

### STM state

This behavioral state represents the *control-capable* state in which mice could exert cognitive control such that their responses were guided by the persistent frontal activity recruited by the stimulus (Supplementary Fig. 11a). The choice under the STM state was thus modeled by the repeat-independent DW (i.e. without any dependence on previous trials). In particular, given a stimulus *S*_*t*_ and a delay *D*_*t*_, the probability to generate a Right response *P*(*R*| *S, D*) was given by the probability that the DW was in the Right attractor at the end of the delay, as given by Eq. 10.

## Fitting of the model

To fit the parameters of the STM and RepL states together with the HMM transition probabilities and the initial state probability Π, we used an Expectation-Maximization algorithm. With this approach we can maximize the log-posterior of the parameter set (Θ) given the trial sequence of stimuli (***S***), delays (***D***) and animal choices (***r***). This approach has already been described in (Ashwood et al. 2022; Baum 1972) and we only adapted it to fit in addition the parameters of the RepL and the STM states. For completeness, we describe the procedure here. Our aim is to maximize the log-posterior of the parameters Θ given the data (***S, D, r***):

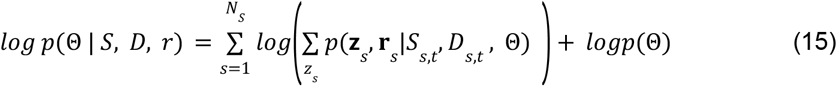

where the first sum is over all experimental sessions (*s* = 1, 2, … N_S_), and the second one is *T* over all 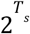 possible latent state paths of session *s* (i.e. all binary sequences of length *T*_*s*_ with elements *z=*{RepL, STM}). To efficiently compute the sum over latent states during the Expectation step, we used a forward and backward pass over the data to compute: 1) the posterior probability of each state (*k* = RepL or STM) for each session (*s*) and each trial (*t*) γ _*s,t,k*_ = *p*(*z*_*s,t*_ = *k*|*S*_*s,t*_, *D*_*s,t*_, *r*_*s,t*_, Θ) and 2) the joint posterior distribution of two of consecutive states (i.e. having states *j* and *k* in trials *t* and *t*+1 of session *s*: ξ _*s,t,j,k*_ = *p*(*z*_*s,t+1*_ = *k, z*_*s,t*_ = *j* | *S*_*s,t*_, *D*_*s,t*_, *r*_*s,t*_, Θ)) (Baum 1972). With these probabilities we can +1write the expected complete data log-likelihood (ECLL), which is a lower bound of the log-posterior of the parameters given the data:

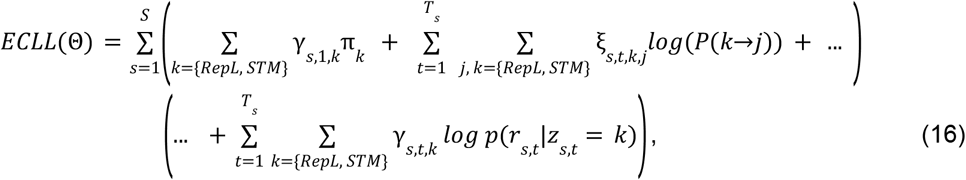

where *P*(*k*→*j*) is the probability to make a transition from the state *k* to *j*, with j, k = {RepL, STM}. During the maximization step, we maximize the ECLL respect to the parameters, something that has been shown to always increase the log-posterior (Bishop 2016; Baum 1972). We iteratively used this algorithm until the change in ECLL was smaller than 10^−5^ log-likelihood units. To fit the DW and the RepL parameters, we used the same Gaussian prior in Eq. 15 that we used for the extended DW (eq. 13) with λ = 0. 01. The transition probabilities P(k→j) and the initial probabilities of each state 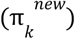, which were not regularized, can be maximized using the classic exact solution developed in (Baum 1972):

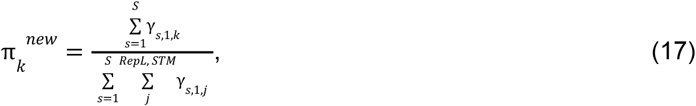

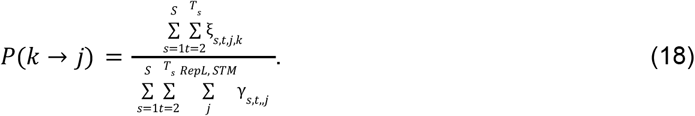

However, we do not have an exact solution to maximize the parameters of the STM state (i.e. the third term in Eq. 16). Thus, to optimize the parameters of both states we used the LBFGS algorithm implemented in the Julia Optim package together with ForwardDiff package to automatically compute the gradients. Similarly to the extended DW model, we ran the algorithm with 100 random initial conditions and we selected the parameters that best fit the data.

### Trial classification

After fitting the parameters of the model, the HMM provides the probability of each of the two behavioral states for each trial *t* in the posterior probability γ _*s,t,k*_of the latent state variable *z*_*t*_ to be either STM or RepL: in the Results section, for simplicity, we rename these probabilities as *p*(*z*_*t*_ = *STM*) and *p*(*z*_*t*_ = *RepL*) = 1 − *p*(*z*_*t*_ = *STM*). To further ease the notation, we used *p*(*STM*) = *p*(*z*_*t*_ = *STM*). We then classified trials either as *STM trials* or as *RepL trials* depending on whether *p*(*STM*) > 0. 5 or *p*(*STM*) < 0. 5, respectively. In some sessions we observed fast excursions of *p*(*z*_*t*_ = *STM*) *t* below 0.5 lasting only one trial. To make the classification more robust, we requested STM and RepL phases to last more than one trial. We imposed this by first performing a running average of *p*(*z*_*t*_ = *STM*) with a 3-trial window and then using this *filtered* probability trace to classify *t* each trial based on the threshold 0.5.

### Neural Population Analysis

After spike sorting, we obtained *N*_*units*_ single units in each session (mean *N*_*units*_ = 2089; range 7-162). From the spike trains of these single units we generated population activity vectors **n**_*k*_ (*t*) for each trial *k*, in which each entry *n*_*i,k*_ (*t*) represented the spike count of the *i*-th neuron in the count bin (*t* − Δ*t*/2, *t* + Δ*t*/2), where the time *t* is taken relative to the Stimulus Cue onset *T*_St_ or to the Go Cue onset *T*_Go_. We used non-overlapping count bins of Δ*t* = 250 ms, except in Figs. 6i and Supplementary Fig. 5 in which we used Δ*t* = 500 to speed up the analysis as the temporal window spanned was very large. We then performed logistic regression to, based on a linear combination of the spike counts *n*_*i,k*_ (*t*) at a given time *t*, classify that trial *k* according to the animal’s choice *r*_*k*_. We defined the linear combination of spikes counts **n**_*k*_ (*t*) with the decoding weights **β** as the log odds of the classification in that trial *x*_*k*_ (*t*) = **β** · **n**_*k*_ (*t*). We account for the offset β _*0*_ in the classification boundary by adding an extra entry *n*_*N*+,*k*_ =1 in the spike counts vector **n**_*k*_ (*t*). The probability that the decoded choice 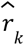 was a Rightwards choice can then be expressed as the log odds passed through a logistic function ϕ(·):

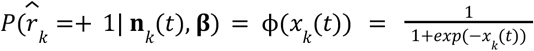

For each recording session, we fitted the vector of weights **β**(*t*), i.e. the decoder, by maximizing the log-likelihood of the animal choices ***r*** counts: given the decoder and the spike counts:

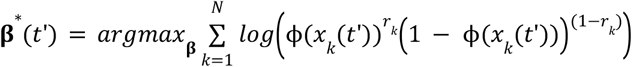

Depending on the training time *t’*, the decoder that maximizes the likelihood is different and hence we include the time dependence. We used cross validation by using different trials for the training of the decoder and for testing its accuracy (see below; Table 2).

**Table 2.**
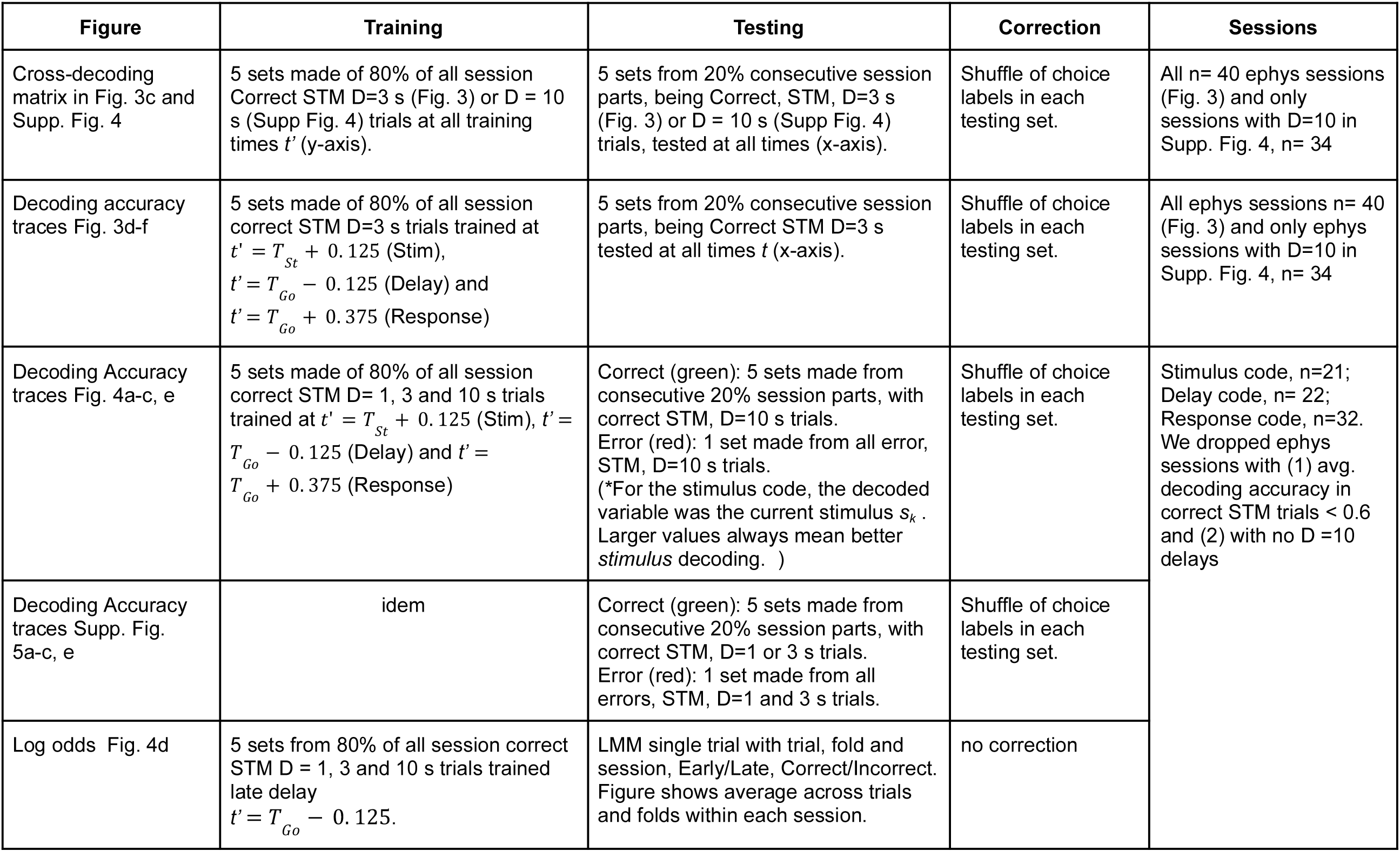
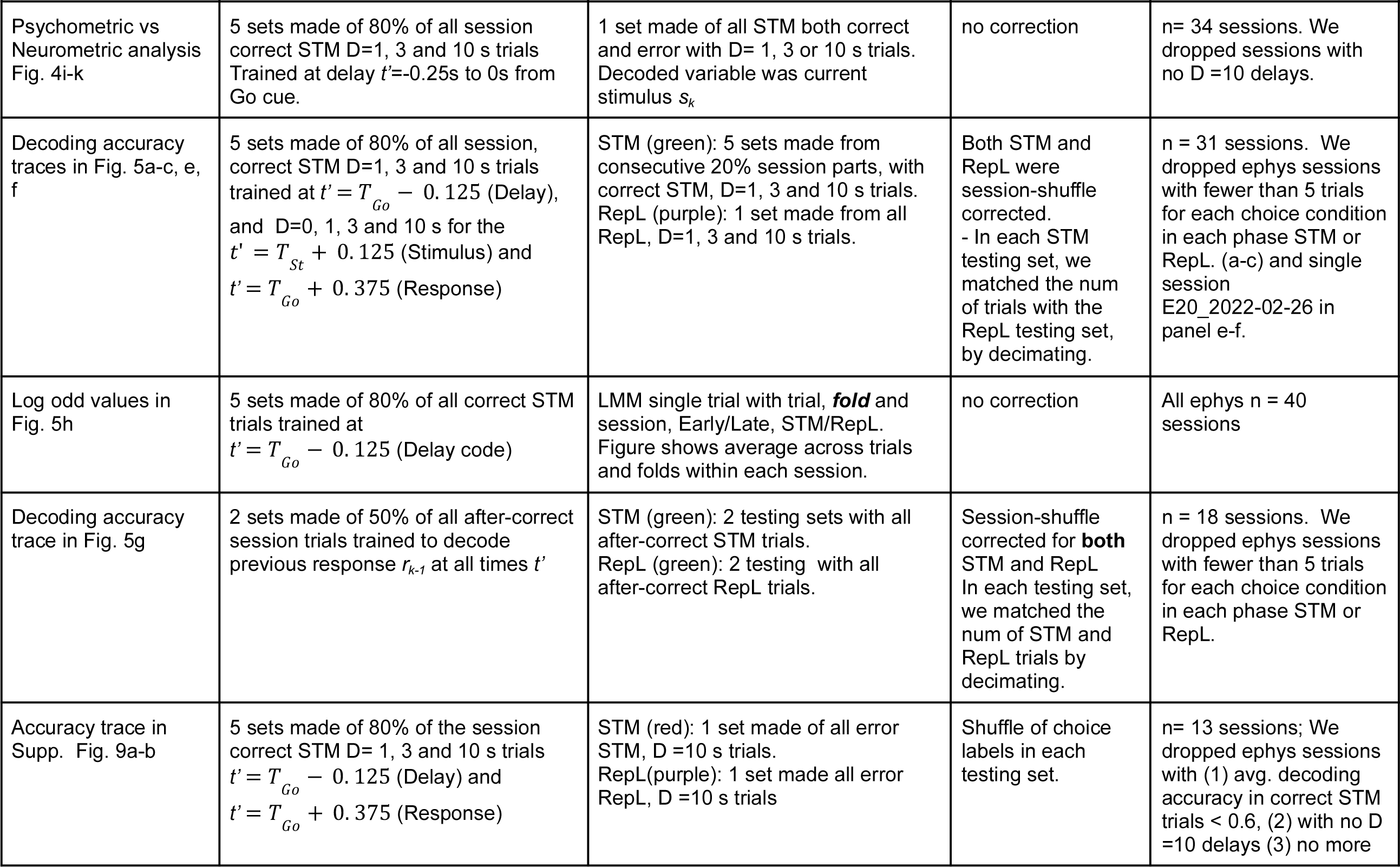
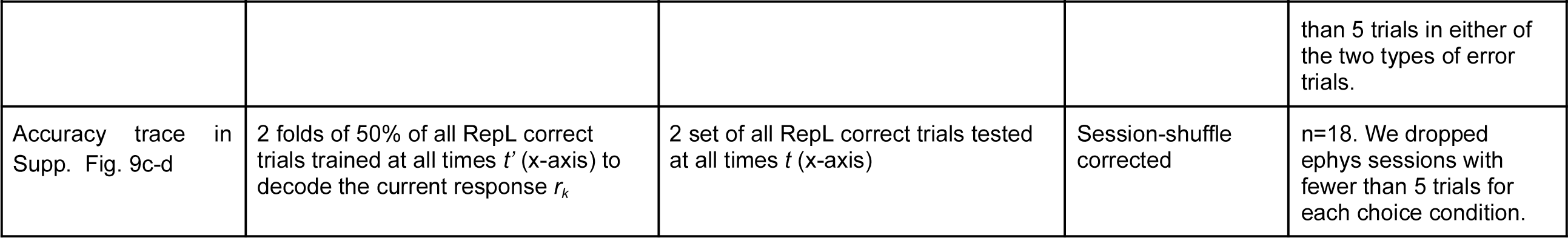
Methodological details about the population decoding analysis for each of the figures of the manuscript.

### Cross-decoding Matrix

The cross decoder’s accuracy *DA*(*t, t*’) shown in Fg. 4c and Supp.

Fig. 4a, shows the probability that the decoder **β** (*t*’)^*^ trained at time *t’* (y-axis) predicted the animal’s choice from the population activity **n**_*k*_ (*t*) at the *testing* time *t* (x-axis):

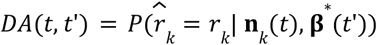

We used 5-fold cross validation on correct trials labeled as STM by the HMM (i.e. 5 testing sets consisting of 20% of consecutive trials from each session, the other 80% trials as training set). The decoding accuracy was then obtained by averaging across folds and then across sessions. For each matrix, we restricted both training and testing to either only D= 3 s (Fig. 3c) or D = 10 s trials (Supp. Fig. 4a).

### Stimulus, Delay and Response STM Decoders

The three decoders presented in Figs. 4, 5 and 6 (except panel i), Stimulus, Delay and Response, differ only on the time in which the the decoder was trained: the *Stimulus code* **β**_*Stim*_ = **β**^*^(*t*’ = *T*_*s,t*_ + Δ*t*/2), the *Delay code* **β** _*Delay*_ = **β**^*^ (*t*’ = *T*_*Go*_ − Δ*t*/2) and the *Response code* **β** _*Resp*_ = **β** ^*^ (*t*’ = *T*_*Go*_ + 3 * Δ*t*/2). All three were trained using correct trials labeled as STM by the HMM. Each code was then tested in four different conditions: STM correct trials (Fig 4 and 5, green lines), STM error trials (Fig. 4 red lines), RepL correct trials (Fig. 5a-h, purple lines) and RepL error trials (Supplementary Fig. 9a-b, red lines). For correct STM trials, because the condition in training and testing was the same, we used 5-fold cross validation. For the other three conditions, training and testing sets contained different trials, so there was no need to cross-validate.

The Stimulus code was defined to predict the current stimulus *S*. Because it was trained in *k* correct trials where the stimulus and animal’s response matched, this distinction between predicting the stimulus instead of the response was only necessary when tested in error trials (Fig. 4a red trace). There, we computed the accuracy that the decoded response matched the current stimulus 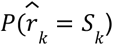, so that larger values in decoding accuracy meant better *stimulus* decoding.

To compare decoders between STM and RepL trials (Figs. 6a-d, h), we downsampled the testing set of STM trials to match the trial number found in RepL for each of the sessions. The training dataset was not downsampled for this. In the case that a given session had fewer than 5 RepL trials for testing in any of the two stimulus conditions, that session was removed for both STM and RepL.

To control for small imbalances in the number of Left and Right choices in a particular session, we subtracted the decoding accuracy obtained when the choice labels in the testing set had been shuffled across trials. We generated 100 of these shuffles of the testing set for every fold, and subtracted the mean decoding accuracy of the shuffles. The remaining accuracy after removing the choice-shuffled accuracy was defined as the Excess in Decoding Accuracy (see Table 2 for details).

Plots showing the temporal course of the excess of decoding accuracy typically show the mean across the sessions that meet the criteria for the analysis (see Table 2 for details) with a 95% confidence band obtained by bootstrapping over sessions (Fig. 3d-f, Fig. 4a-c and Fig. 5a-c, g)

### Psychometric versus Neurometric analysis

We computed neurometric accuracy curves using the Delay code **β** _*Delay*_ to predict the current stimulus at every time point during the delay using both correct and error trials. In each session, we computed the neurometric accuracy by signing the log odds of the neural classification *x*_*k*_ (*t*) = **β** _*Delay*_ ***n***_*k*_ (*t*) according to the stimulus: *x*’ _*k*_ (*t*) = *x*_*k*_ (*t*) for Right stimulus trials and *x*’ _*k*_ (*t*) =− *x*_*k*_ (*t*) for Left stimulus trials. The neurometric accuracy in log odd units was then the trial average 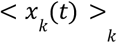 (Fig. 4i-j, gray traces) and the neurometric forgetting rate as the slope of a linear fit in the interval 0 < *t* < 10 s (Fig. 4i-j, blue line). We then compared the neurometric forgetting rate with psychometric forgetting rate equivalently defined as the slope of the linear fit of the log odds *y*_*k*_ (*D*) of the behavioral choice accuracy, i.e. probability *p* (*D*) that the mouse responded correctly at delay *D =* 1, 3 and 10 s, averaged across trials (Fig. 4i-j, orange line 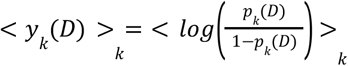 We used only n=34 sessions that included the longest *D* =10 s (Fig. 4k) as with shorter delays the slope of the curve was too noisy to be estimated reliably (see sessions in Supplementary Fig. 8. with only *D =* 1 and 3 s).

### Decoding of previous response

We wondered whether the tendency to repeat the previous choice was greater in RepL trials because there was a stronger neural trace of the previous choice bridging the previous and current trial (Barbosa et al. 2020). To test this, we quantified the extent to which the previous response could be decoded during the following inter-trial-interval and during the current trial and compared STM versus RepL trials (Fig. 5g). Specifically, we trained a family of decoders to classify each trial according to the previous choice, based on the spike counts at each time point. We spanned the interval from previous response onset to the current stimulus onset and until to current Go cue onset (Fig. 5g). We used all STM and RepL trials following a correct choice to avoid the Timeout interval caused by an error in the previous trial. Because the number of RepL trials was small in some sessions, we used 2-fold cross-validation, creating a training set with 50% of trials with both STM and RepL and then testing separately the STM and RepL trials in the other 50% testing set.

### Session shuffle analysis

Since choices within the RepL state had a strong repeating tendency, they strongly violate the assumption of independence between trials implicit in most statistical tests. In particular, it has been recently shown that when behavioral variables show strong serial correlations because they either show slow across-trial fluctuations, they follow a trial block design or simply because they exhibit nonstationarities across the session, an accurate characterization of chance levels using adequate permutation tests is necessary to avoid spurious correlations (Harris 2020; Elber-Dorozko and Loewenstein 2018). For this reason, in the analyses that included the decoding accuracy in RepL trials (Fig. 5a-c,e-g; Supplementary Fig. 9a-d; see Table 2 for details), instead of the trial label shuffle used for the other decoders, we performed a session shuffle decoder correction (Harris 2020). For this, we took the recordings from each electrophysiology session and paired them with the sequence of choices from another randomly chosen behavioral session of the same mouse. Then we trained the neural decoder to predict those choice labels using the neural activity exactly in the same way as with the non-shuffled data (i.e. using cross-validation). The paired behavioral session had to have similar properties as the electrophysiology session, in particular it ought to have at least as many trials in each of the STM and RepL phases as the electrophysiology session. The excess of trials in each phase of the behavioral session were removed to exactly match the number of trials of the electrophysiology session. For each electrophysiology session, we trained and tested this session-shuffled decoder with all the behavioral sessions that met those criteria and then averaged the resulting accuracies. This average was taken as the chance level baseline and was subtracted from the decoding accuracy of the non-shuffled decoder yielding the Excess of decoding accuracy we finally plotted. In Supplementary Fig. 9d-e we show this spurious above-chance decoding accuracy found in RepL trials when the session-shuffle correction was not applied.

### Overlap between population codes

To quantify the overlap between codes, we computed, the pairwise cosine of the angle between the code vectors **β** _*Delay*_, **β** _*Stim*_ and **β** _*Response*_.

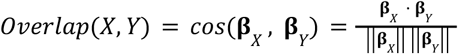

To quantify the stability of the Delay code, we distinguished between two different Delays codes depending on when the code was trained: the Late Delay code **β** _*Late*−*Delay*_ = **β** _*Delay*_ was defined at the end of the delay period as described above. The Early Delay code was computed as **β** _*Early*−*Delay*_ = **β**^*^(*t*’ = *T* _*Stim*_ + 500 *ms* + Δ*t*/2). Values of the overlap close to 0 represent independent codes whereas overlap close to 1 refer to strongly overlapping codes. However, because our estimate of each code is subject to statistical error given by a finite number of trials, in practice it is difficult to reach overlap values near 1. To have a reference of what the overlap should be between two “almost identical” codes, we defined a third Late’ Delay code **β**_*Late*’−*Delay*_ = **β** ^*^ (*t*’ = *T* _*Go*_ − 3Δ*t*/2), which was trained in the bin Δ*t* just before the Late Delay code (Fig. 3c, vertical bars on the right). We thus interpreted the overlap between Late and Late’ as the maximum overlap we could obtain between two codes (Fig. 3g)

### Analysis of Brain State

To quantify brain state changes across the recording sessions, we focused on population synchrony. We only used sessions with a number *N* of well isolated units *N* ≥ 30 because having fewer simultaneously recorded units compromises the quantification of population synchrony. We hence used n=31 electrophysiology sessions out of a total of 40 recorded from 8 mice. We estimated the degree of synchrony among neurons in the population from the variance of the population instantaneous spiking rate (Mochol et al 2015; Renato et al 2023). The population instantaneous rate in the *k*-th trial of a given session was defined as 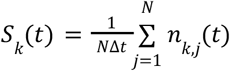 where the sum goes over the spike counts of all the *N* well isolated units of the session. To assess brain state we used spike count bins of Δ*t* = 20 ms. For each trial, we computed the standard deviation of the spike count in these bins across time σ_*k*_ = *StdDev*[*S* _*k*_ (*t*)]. We used the pre-stimulus interval defined as the 2 s period just before the stimulus onset. We normalized the standard deviation in each trial by the standard deviation 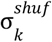 that would be obtained if neurons were asynchronous homogeneous Poisson processes. We computed 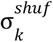 by creating 100 shuffled surrogates from *S* _*k*_ (*t*) where the spike times were randomly shuffled. We defined a trial-by-trial pre-stimulus synchrony index as the ratio between the two standard deviations:

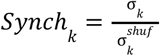

The resulting synch coefficient *Synch* is close to 1 when the neural activity is asynchronous, *k* whereas higher values indicate the degree of population synchrony (Fig. 6b-c). We computed the across-trial correlation coefficient of *Synch* with the *a posteriori* probability of the HMM to be in STM *p*(*z*_*k*_ = *STM*) (Fig. 6d): *CorrCoef* [*Synch* _*k*_, *p*(*z*_*k*_ = *STM*)]. To correct for correlation arising from the non-stationarity of the *Synch*_*k*_ and *p*(*z*_*k*_ = *STM*) traces across the session (Supplementary Fig. 10), we computed the correlation coefficient after pairing the *Synch*_*k*_ trace from each session with the behavioral trace 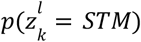 from a different behavioral session *l* randomly chosen from the same animal (Harris 2020). For each recording session, we paired its *Synch*_*k*_ with all behavioral sessions with the same or more trials, and computed the average correlation coefficient across session-shuffled pairings. The corrected correlation coefficient of a given session was then 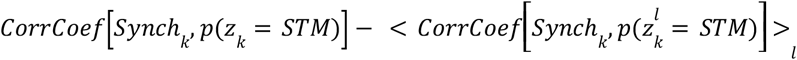 The same method was applied to compute the correlation of the synchrony index *Synch*_*k*_ with (1) the binary *k* sequence of choice outcomes *o*_*k*_ (with *o*_*k*_ = 1 for correct trials and zero otherwise); (2) the binary sequence of repetitions *rep*_*k*_ (with *rep*_*k*_ = 1 for trials with a repeated response *r*_*k*_ = *r*_*k*−*1*_, and zero for an alternating response) (Fig. 6d). The significance of the corrected correlation coefficients from all sessions was assessed using a two-tailed *t* test (Fig. 6d),

To assess the degree of rhythmicity of the population activity during periods of synchronization, we computed population instantaneous rates *S*_*k*_ (*t*) using finer bins of Δ*t* = 5 ms. We computed the autocorrelogram of *S*_*k*_ (*t*) in each trial and then averaged them, *k* separately for STM or RepL trials in each session (see example session in Fig. 6e). Autocorrelograms were not normalized by the variance but measured the raw covariance between the population rate and its lagged copy yielding units of rate squared. The value at time lag=0 was removed for visualization. Comparing the average autocorrelogram of RepL versus STM trials, revealed an enhanced oscillatory structure in RepL trials of many sessions (see all individual sessions in Supplementary Fig. S13). Because the frequencies were slightly different between sessions and in some sessions there were no traces of oscillatory activity, the session-averaged auto-correlogram in RepL trials did not exhibit oscillatory traces as clearly as individual sessions. We then computed the power spectral density (PSD) of

*S* (*t*) *k* using Welch’s overlapped segment averaging estimator. The session-averaged PSD for RepL and STM trials revealed a clear increase in power in RepL trials in the band ∼ 3 − 15 Hz (Fig. 6f). To obtain a more quantitative measure of the peak frequency, we computed the ratio of PSDs and found that the increase in power peaked at 4.4 Hz (Fig. 6f right). We also computed the PSD ratio for each session separately and computed the highest peak in the range [2, 15] Hz (see red dots in Supplementary Fig S13). The median of the peak frequencies was 4.91 Hz with Q1-Q3 quartiles 4.06 - 7.88 Hz (histogram in Supplementary Fig S13).

