## Supplemental Figures for "Episodic recruitment of attractor dynamics in frontal cortex reveals distinct mechanisms for forgetting and lack of cognitive control in short-term memory"

### Supplementary Figures.

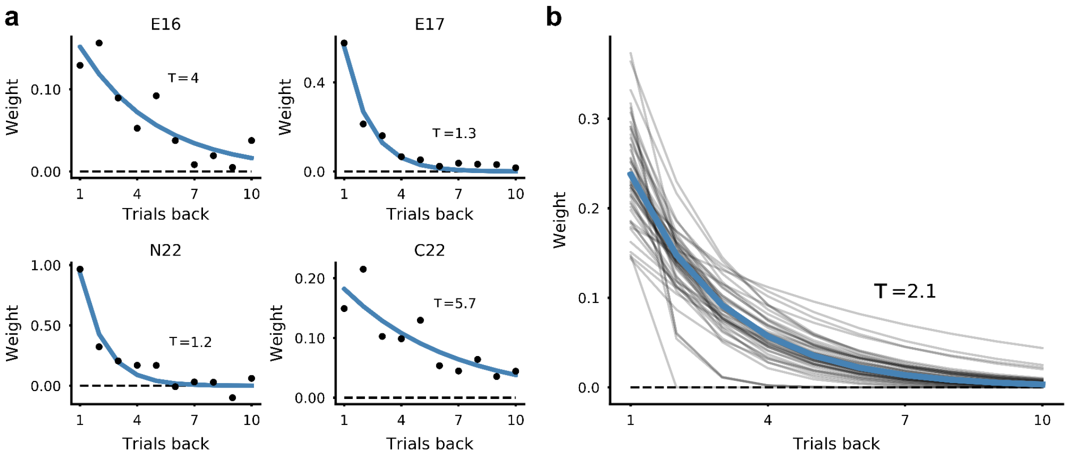


**Supplementary Fig. 1. Impact of previous choices on the current choice. a,** GLMM fitted weights of previous choices as a function of the number of trials back into the past (dots) for four example mice (see panel titles). Weights were obtained from fitting the current choice using the GLMM shown in Fig. 1k except for the *Action trace* regressor $C_t$ which was replaced by the ten previous choices whose impact was assessed independently. Positive values of the weights imply a tendency to repeat previous choices. The outcome of previous choices was not taken into account. Exponential curves were fitted to the weights and their time constant τ is shown for each animal. **b,** Exponential fits from all the animals (Groups #1 and #2, n=48). Thick blue line represents the mean curve with its corresponding τ (inset).

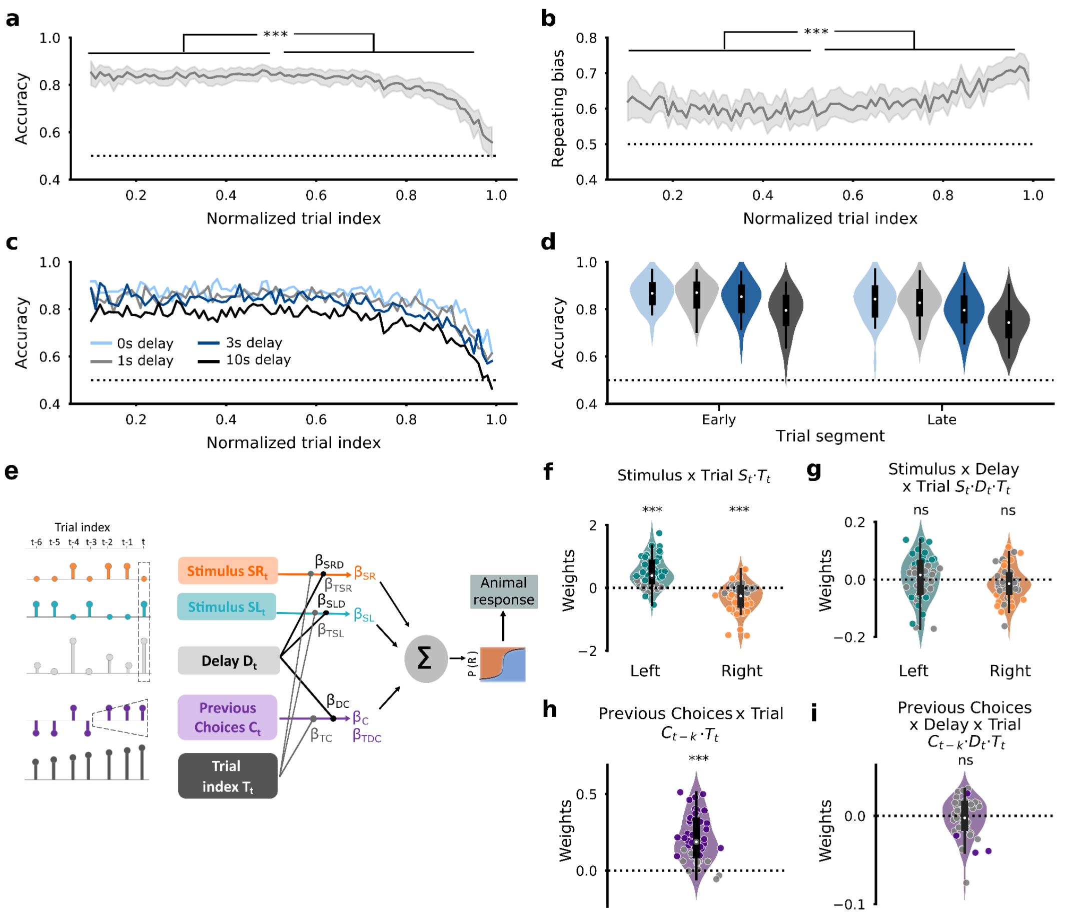


**Supplementary Fig. 2. Effect of trial index on mouse choice behavior. a-b**, Averaged accuracy and repeating bias versus the within-session normalized trial index *T,* where $T = 0$ and $T = 1$ represent the beginning and the end of each session. *Early* in the session was defined as trials $0 < T \leq 0.5$ and *Late* as $0.5 < T \leq 1$. t-test for paired samples with Bonferroni correction **** $p<10^{-4}$. Group #2, n=34 (panels a-d) **c**, Accuracy versus normalized trial index separately computed for each delay length (inset). **d,** Accuracy versus delay length separately computed for Early and Late trials. **e**, GLMM schematic extending the one shown in Fig. 1k by including the interactions of the normalized trial index *T* with the Stimulus, the Delay and the Prev. Choices (see Methods). **f-i.** GLMM fitted weights showing the interaction between stimulus x *T* (f), between stimulus x delay x *T* (g), Action trace x *T* (h) and Action trace x delay x *T* (i). Dots represent individual mice (Groups #1 and #2, n=48). Dot color represents individual significance ($p < 0.05$). While the increase of trial index diminishes the impact of the stimulus (f) and increases the impact of previous choices (h), it does not alter their dependence on delay (g, i). This implies that the stability of the memory (captured by Stim x Delay, Fig. 1m) and the absence of delay dependence in the Repeating bias (Fig. 1o)) do not co-vary with the drift in performance across the session (a).

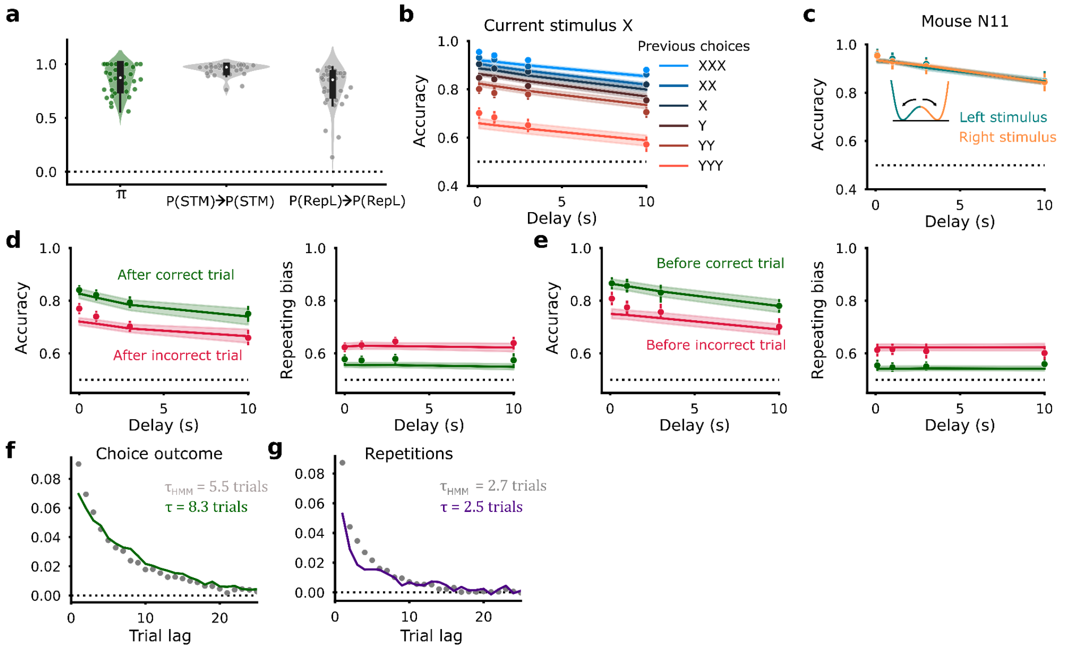


**Supplementary Fig. 3. Results from fitting the behavioral HMM and comparison with mouse data. a**, Fitted parameters of the HMM model: $\pi = p(z_1 = STM)$ (probability of starting the session in the STM state), P(STM→STM) (probability of staying in the STM state) and P(RepL→RepL) (probability of staying in the RepL state). Each dot represents one animal (Group #2, n=34). **b**, Mean choice accuracy versus delay conditioned on whether the previous choices were the same (X) or the opposite (Y) to the current stimulus side. Conditioning on longer streaks of previous choices is indicated as X, XX and XXX for one, two or three same-side choices and similarly for opposite side choices (Y, YY and YYY). Dots indicate mouse averaged behavioral data whereas lines show the accuracy given by the fitted HMM. **c**, Accuracy versus delay computed separately for Left (green) and Right stimulus trials (orange) for an example mouse showing an unbiased behavior (dots) that was capture in the HMM (lines) by having equal initial loading probabilities $P_L$ and $P_R$ and a near zero asymmetry parameter $\mu$ (see double-well cartoon). **d-e,** Mean choice accuracy (left) and repeating bias vs. delay (right) conditioned on whether the previous trial (d) or the following trial (e) was correct (green) or incorrect (red). Dots correspond to average behavioral data and lines to the HMM. The lower accuracy and larger repeating bias after incorrect trials could erroneously be interpreted as mice tending to repeat more after error choices. However, the difference is also observed when conditioning on the *next* trial and it can be reproduced by the HMM which does not take into account previous outcomes (the HMM only considers previous choices). The result can be simply explained by the difference in accuracy between STM and RepL phases: conditioning on either the previous or the next trial being incorrect, biases the trial sampling towards the RepL phases which overall have lower accuracy and larger repeating bias. **f**, Average autocorrelogram of choice outcome sequences (f) and repetition sequences (g) obtained from the mouse data (dots) and from the average fitted HMM (lines). Data autocorrelograms were corrected to remove the influence of within-session drifts (Methods). Inset shows the values of the time constant $\tau$ obtained from exponential fits to either the mouse of the HMM data.

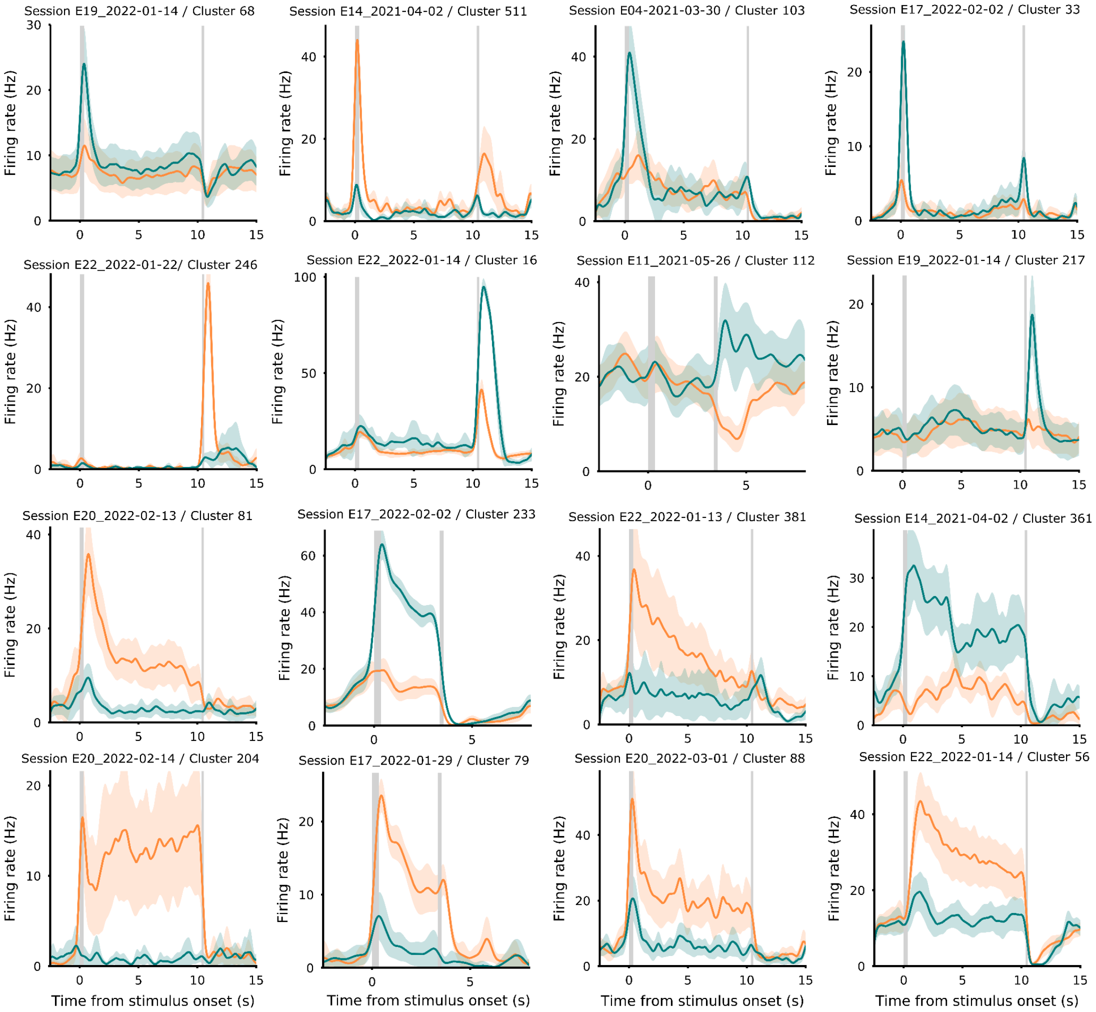


**Supplementary Fig. 4. Average trial activity for some example neurons.** Each panel shows the firing rate of a single cell averaged across correct STM-classified Right (orange) and Left stimulus trials (teal) with one of the three delays (i.e. either D = 1,3 or 10 s). For each trial, a rate trace was obtained by convolving spike trains with a Gaussian kernel ($\sigma$ = 200 ms). Plots depict mean traces across trials, and error bars indicate 95% CI. Vertical lines represent the stimulus and the Go cue. While many neurons were only selective in their response to the stimulus or in their activity during the licking action, a substantial fraction of neurons showed persistent selective activity.

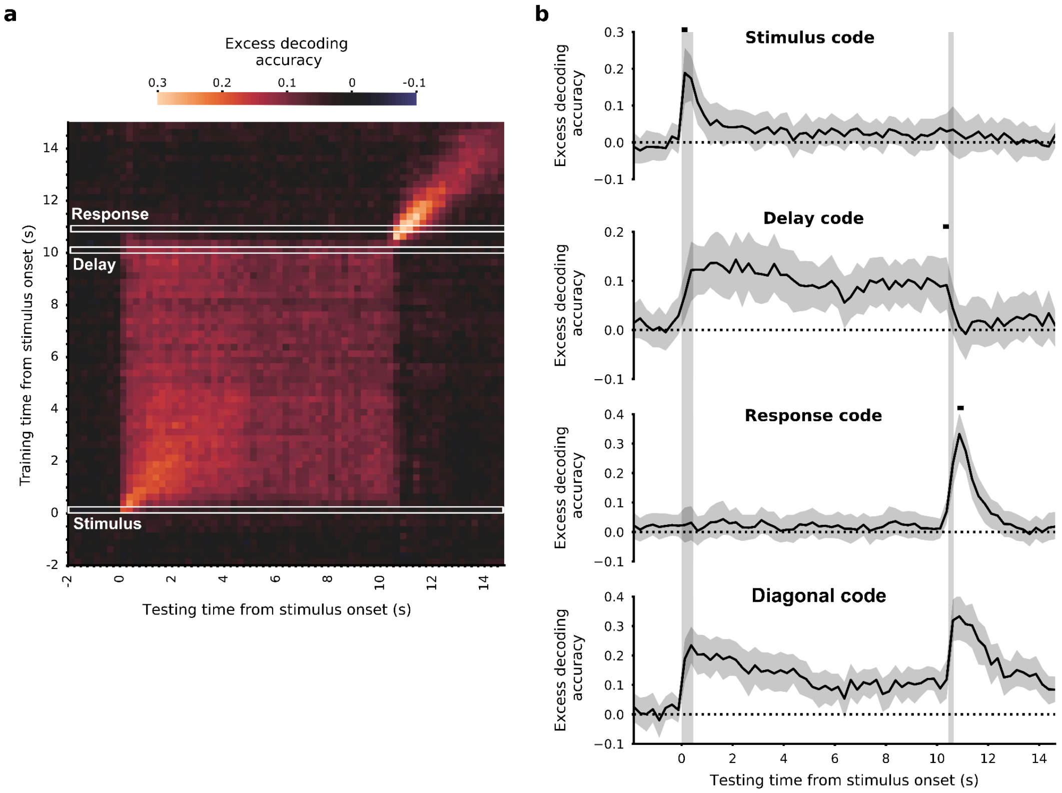


**Supplementary Fig. 5. Dynamics of ALM population codes across 10 s delay trials. a,** Average cross-validated excess of decoding accuracy as a function of when the decoder was trained and tested to predict the animal's choice. Training and testing was done with correct 10s trials 5-fold cross-validated (n=34 sessions; spike count bin size was $\Delta t$ = 500 ms). The Stimulus code, Delay code and Response code are defined by the training time intervals (0, 0.25) s, (10, 10.25) s and (10.75, 11) s (white horizontal rectangles). Diagonal code (not marked) represents the diagonal of this cross-decoding matrix, with training and testing performed in the same bin. **b**-**e,** Time evolution of the average cross-validated excess of decoding accuracy of the Stimulus, Delay, Response and Diagonal codes. The time axes of all plots were aligned to stimulus onset. Small black marks above each trace indicate the time interval used for training each code (except for the diagonal code that was trained in each time bin). Error bars in b represent 95% confidence intervals obtained through bootstrapping of the sessions.

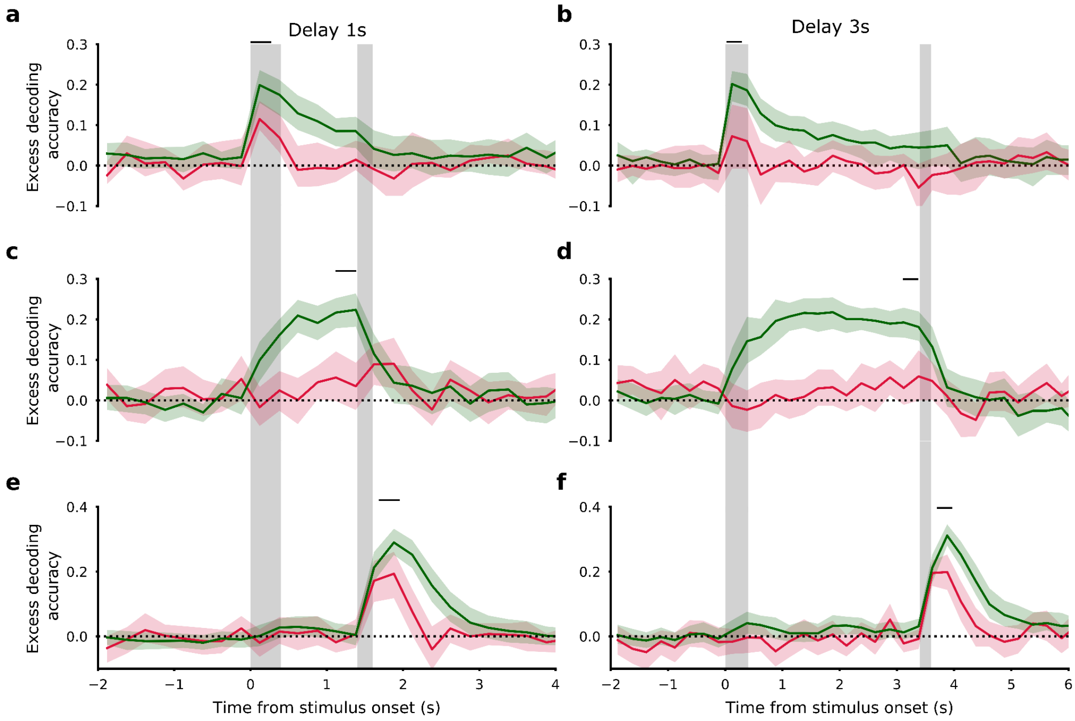


**Supplementary Fig. 6. Decoding accuracy of population codes during short delays of 1 and 3 s. a-f,** Average cross-validated excess of decoding accuracy of the Stimulus code (a-b), the Delay code (c-d) and the Response code (e-f) trained using all STM correct trials and tested separately in both STM correct (green) and incorrect (red) trials with D = 1s (left panels) and D = 3s delay trials (right panels) (n=40 sessions). Error bars indicate 95% confidence intervals. Vertical bands represent stimulus and Go Cue. Above zero excess accuracy indicates proper stimulus decoding for the Stimulus code and proper response decoding for the Delay and Response codes. Stimulus and Response codes show similar curves in 1s and 3s delay to those found in 10s trials (Fig. 4a,b). In contrast, the Delay code in error trials showed no traces (D = 1 s) or very weak traces (D = 3 s) of memory reversals compared with 10s delay trials (Fig. 4c), as the accuracy after stimulus onset in error trials remains at chance indicating that no memory was loaded in STM during these trials. This suggests that errors in STM trials are caused by at least another factor other than maintenance (e.g. loading). By conditioning on different delay lengths, errors can be dominated by non-maintenance errors (short delays) or by the maintenance errors (long delays). This dependence of the reversal amplitude on the delay is reproduced by our DW model (Supplementary Fig. 7).

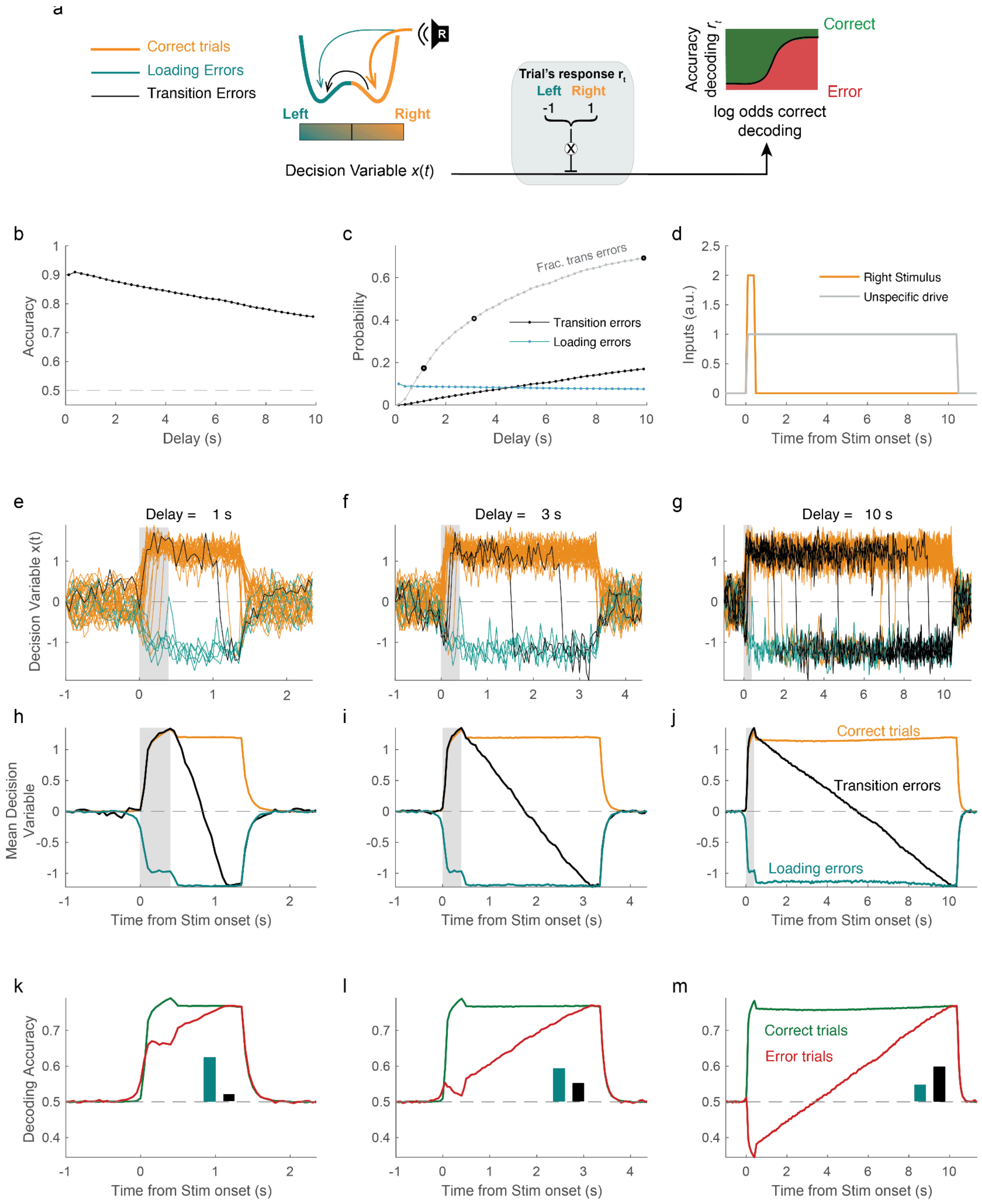


**Supplementary Fig. 7. Performance of the DW model as a function of delay length.**

**a,** Schematic showing the transformation from the decision variable *x*(t) of the DW model (Eq. 5) to the probability of correctly decoding the final response *r* of the model, which comes down to a sign multiplication depending on whether the response is Right or Left and a sigmoidal function (right panel). For the rest of the figure, we use a Right stimulus and define (1) correct response trials (orange) as those in which *x*(t) is positive at the end of the delay; (2) *loading errors* (teal) as those errors (negative *x*(t) at the delay end) in which the initial loading was incorrect (negative *x*(t) at the delay onset) and (3) *transition errors* (black) as those trials in which the initial loading was correct (positive *x*(t) at the delay onset) but, because of a transition from the correct to the error attractor, *x*(t) was negative at the end of the delay. **b,** Accuracy versus delay length obtained from simulations of the DW. **c,** probability to observe a STM *loading error* (teal), or a transition error (red), as a function of delay duration. Fraction of transition errors over the total number of errors is also shown (gray). **d,** temporal profile of the stimulus input (orange) and unspecified external drive which causes the model to exhibit two attractors (gray). **e-g,** example traces of the decision variable *x*(t) (40 trials) for delays 1, 3 and 10 s (see panel titles). In each plot, the number of trials for each type (correct, loading error, trans. error) was chosen according to its probability. Only when the delay is sufficiently long, there are a substantial number of transition errors. Notice that the time axis in each plot covers a different range. **h-j,** average decision variable traces $\overline{x(t)}$ separately computed for correct, loading error and transition error trials at each delay length. **k-m,** Average decoding accuracy traces obtained from transforming the model's decision variable traces (panel a), and separately averaging correct (green) and error trials (red). These traces are to be compared with the decoding accuracy of the Delay code found in the data (for D=10 s shown in Fig. 4c and for D=1 and 3 s shown in Supplementary Fig. 6c-d). Inset bars indicate the proportion of loading errors and transition errors making up the total pool of error trials. Because only for the longest delay D = 10s, there is a sufficient fraction of transition errors, the average decoding accuracy in error trials only shows traces of memory reversals for the longest delay, as observed in the data. To simulate the DW we numerically solved equation 5 using $\boldsymbol{\alpha}$ = 4, $\boldsymbol{\tau}$ = 100 ms, $\boldsymbol{\sigma}$ = 0.88 during the stimulus and delay periods but 0.5 elsewhere, stimulus amplitude $\boldsymbol{\mu}_s$ = ± 2, stimulus duration 400 ms. The response in each trial was decided based on the sign of the average of *x*(*t*) over the last 250 ms of the delay.

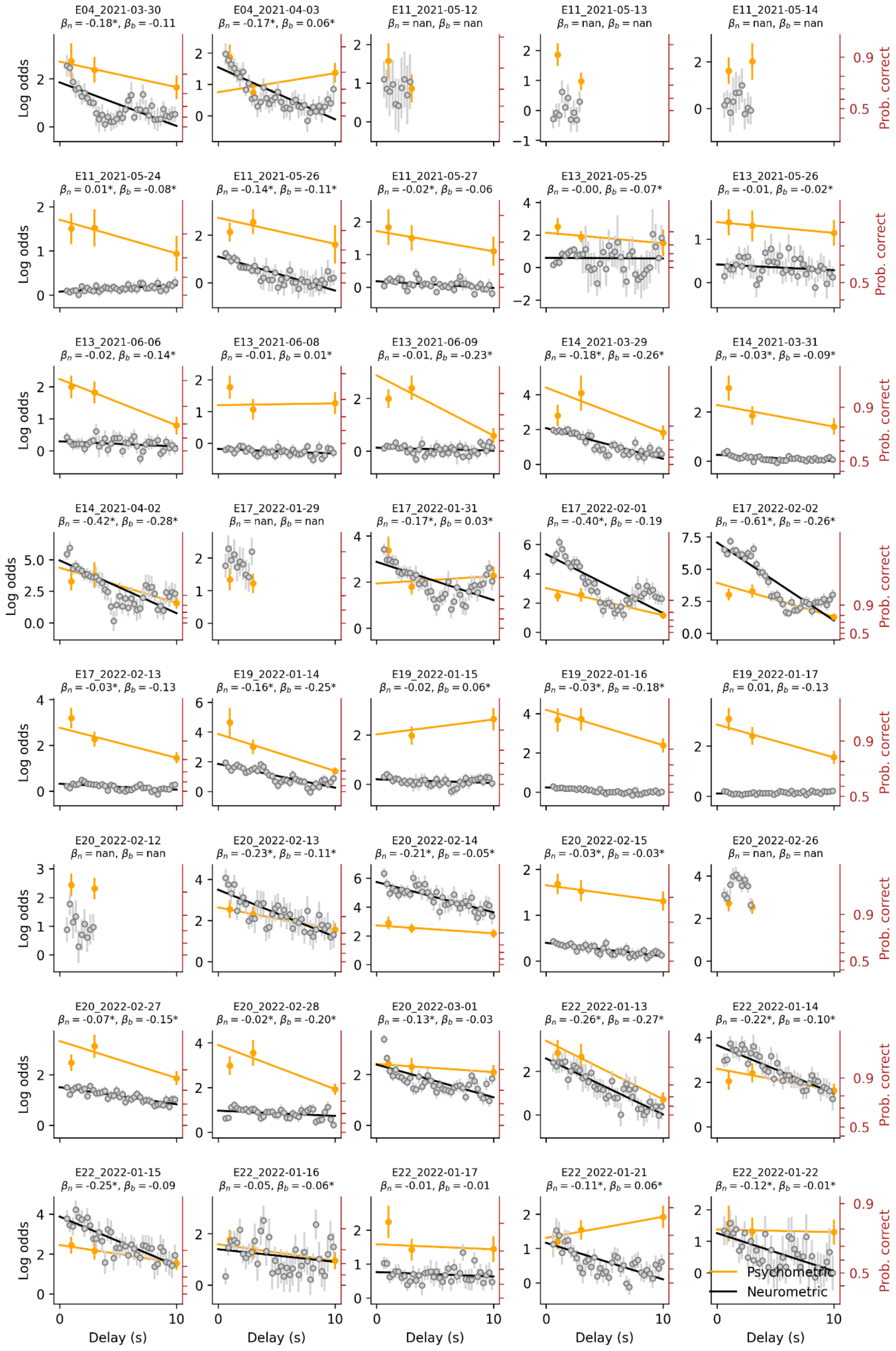

**Supplementary Fig. 8. Comparison of psychometric and neurometric accuracy in each recording session.** For each recording session, each panel shows (1) the probability that the mouse makes a correct response as a function of the delay (orange dots); and (2) the probability of reading out from the ALM population activity the correct response using the delay code at each time point during the delay (gray dots). Probabilities can be read in log odds units (left axes) and in probability units (right red axis). Straight lines are the corresponding linear fits. Titles in each panel indicate the session label and the values of the neurometric slope $\boldsymbol{\beta}_n$ and psychometric slope $\boldsymbol{\beta}_b$ (asterisks indicate $p<0.05$). Error bars show sem. There were 6 sessions out of 40 in which only delays D=0, 1 and 3 s were used (aiming to lower the difficulty of the task that day). These sessions, shown here, were not included in the analysis comparing $\boldsymbol{\beta}_n$ and $\boldsymbol{\beta}_b$ as the decay of the log odds with delay could not be accurately estimated. In session E19_2022-01-15 (4th row, 5th column), all the 53 trials with D =1 s were correct giving an infinite behavioral log odds. Those trials were nonetheless considered for the logit fit of the response outcomes versus delay.

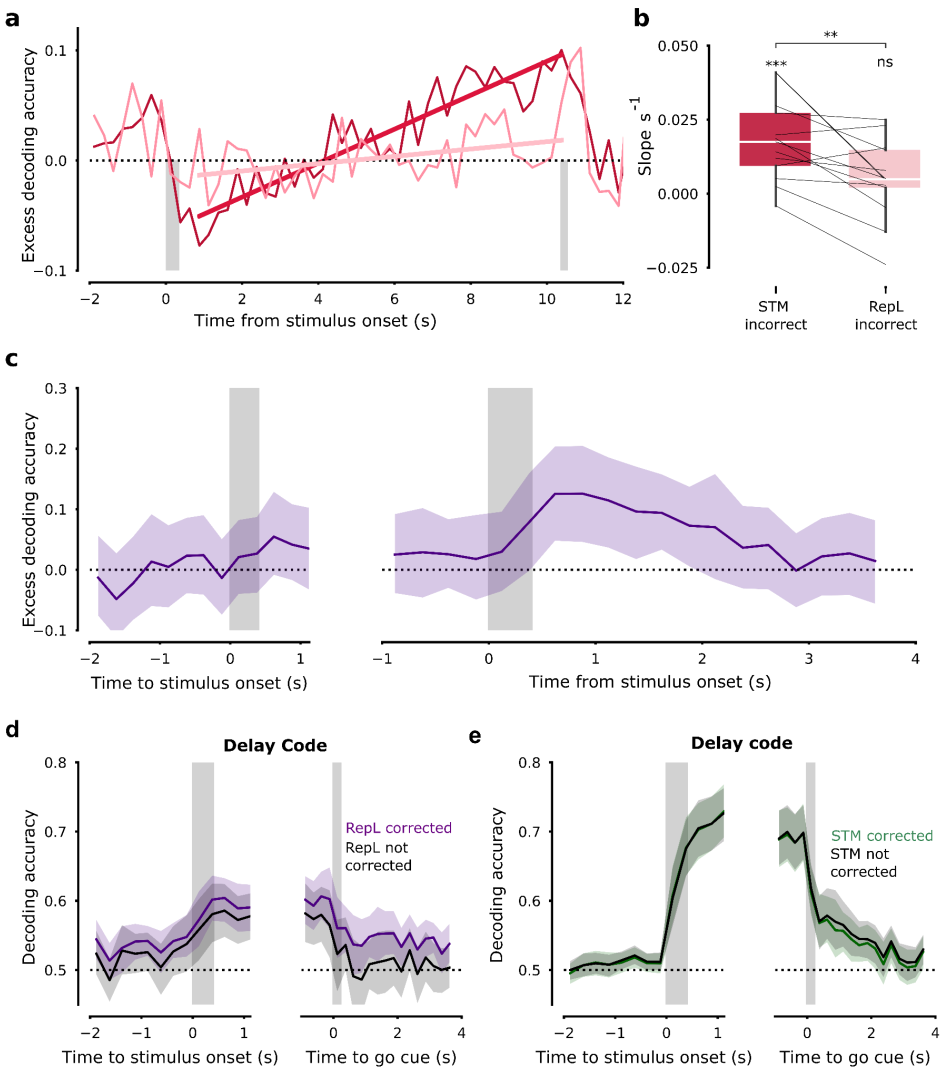


**Supplementary Fig. 9. Comparison of the Delay decoding accuracy in error trials occurring during STM and RepL phases.** Average cross-validated excess of decoding accuracy of the Delay code during error 10s-delay STM trials (red; same trace as in panel Fig. 4c) and during error 10s-delay RepL trials (pink) (n= 13 sessions). The Delay decoder was trained using all correct STM trials and tested in error trials separately for STM and RepL. Positive values of the accuracy indicate decoding of the response opposite to the stimulus (i.e. the incorrect response the animals did in these error trials) whereas negative values indicate decoding of the response associated with the stimulus (i.e. the correct response). Gray shaded areas represent stimulus and Go cue presentation. Straight lines correspond to linear fits of the accuracy during the delay. **b,** paired comparison across sessions of the slope of the Delay code during the delay in error D=10s trials separately taken from STM and RepL phases (i.e. slope of straight lines in panel a, obtained from single sessions). T test against 0 for STM: $t$ = 4.676, $p$ < 5.35 x $10^{-4}$; RepL: $t$ = 1.315, $p$ = 0.21. Two-tailed paired t-test: $t$ = 3.493, $p$ = 4.439 x $10^{-3}$. **c,** Average excess of decoding accuracy of an *alternative decoder* trained using only correct RepL trials. Training and testing were done at the same time (i.e. the diagonal of the cross-decoder). No significant decoding of the final response can be found in the prestimulus, after the stimulus or during the delay and only a Response code seems to be identified. This suggests that ALM has no alternative code at play during RepL trials that, equivalently to the Delay code in STM trials, can sustainably encode the upcoming response before the Go cue. **d-e,** Non-corrected decoding accuracy of the Delay code tested in RepL (d) and STM (e) correct trials. Because choices in RepL phases show strong sequential correlations, decoding accuracy in these trials shows a residual, above-chance decoding baseline which is removed by the session-shuffled correction (compare with the corrected purple trace in Fig. 5c; Methods). This residual decoding accuracy was not present in STM trials. Shaded band in panels c-e represents 95% C.I.

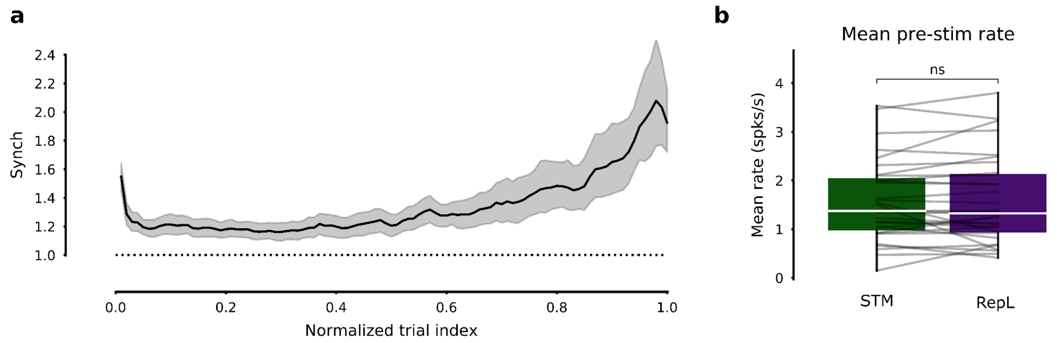


**Supplementary Fig. 10. Brain state changes systematically across the session. a,** Synchrony index versus normalized trial index averaged across sessions (n=31 sessions, error bars are 95% ci). Trial index was normalized by the maximum trial index in each session providing a value between 0 and 1. Synch, computed from pre-stimulus activity (period (-2, 0) from stimulus onset), increases systematically across the sessions, particularly during the second half. **b,** Mean pre-stimulus firing rate for STM and RepL trials shows no significant difference. Each pair of values represents one session. Mean rate in each session was obtained by averaging across STM and RepL trials separately for each unit, and then averaging across all neurons recorded in the session.

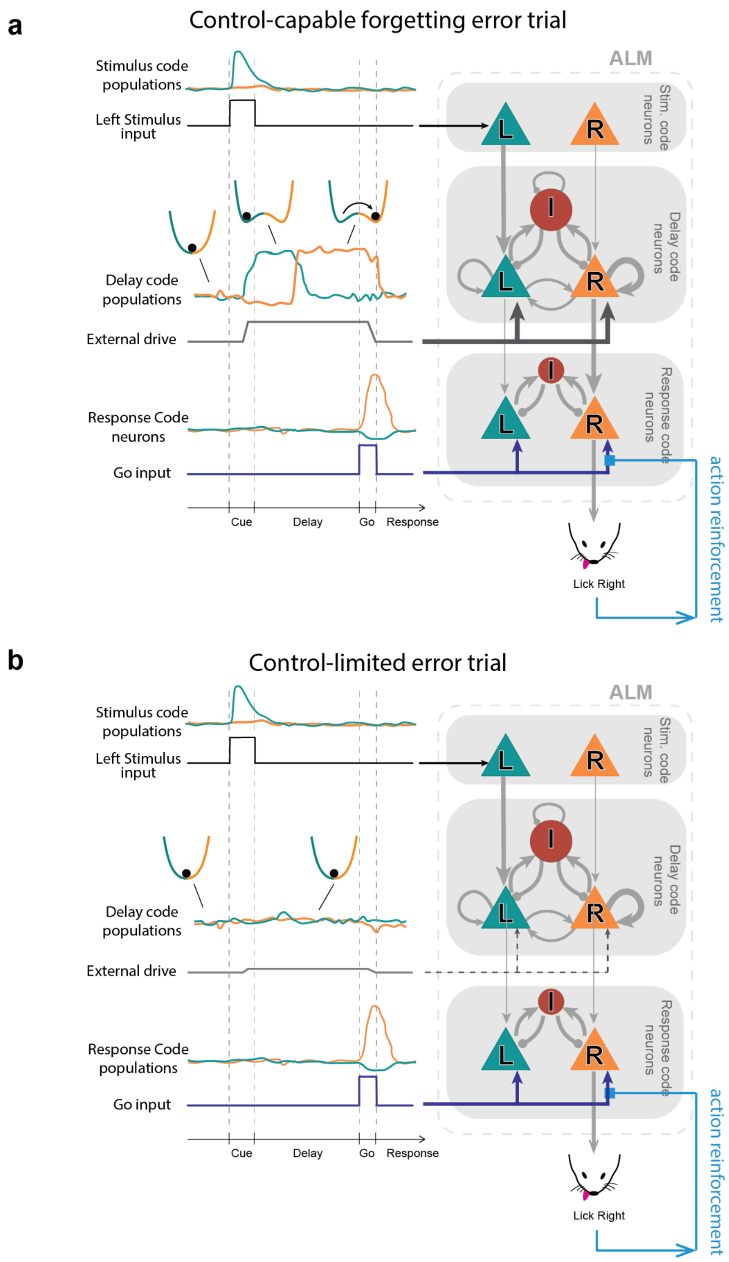


**Supplementary Fig. 11. Schematic of ALM network dynamics during different types of error trials. a,** Proposed dynamics of frontal area ALM during a control-capable forgetting error trial. In trials in which the animal exerts cognitive control, Stimulus code neurons encoding Left and Right choices (top ALM box), receive an excitatory, stimulus-specific input (black upper trace) which makes the corresponding population respond transiently (teal upper trace). At the same time, the Delay code populations (middle ALM box) receive an excitatory external drive (gray middle trace) that remains active during the entire delay. This drive allows the dynamics along the code defined by these two populations to switch from a mono-stable regime in which both Left and Right population fire at the same basal rate (illustrated by a single well potential) to the winner-take-all regime described by double-well attractor dynamics, where it remains during the entire delay (Prat-Ortega et al. 2021; Inagaki et al. 2019). Because of the direct connections between Stimulus and Delay populations, the stimulus information is correctly loaded into the Left attractor generating persistent activity in the

Left-choice-encoding Delay neurons (teal middle trace). During the delay however, maintenance fails and there is a reversal of the system from the Left to the Right attractor so that at the end of the delay, Right-choice-encoding delay neurons exhibit persistent activity. The Response populations are composed of Pyramidal tract neurons that project to motor centers in the midbrain (Inagaki et al. 2022; Economo et al. 2018) ultimately driving Left or Right licking. The presentation of the Go cue provides an excitatory input from the thalamus (blue bottom trace) (Inagaki et al. 2022) to these two populations which, via some inhibition-mediated winner-take-all dynamics, compete to drive the animal's response. The Go cue provides comparable input to the two populations and hence does not provide a decisive signal to unbalance the activity of the Response populations in one way or another. It is the topographic input from the Delay into the Response populations, which, if there is sustained activity at the time of the Go cue, provides a strong input to the Right licking population, which biases the competition and dictates the final response of the animal. **b,** In a trial with no cognitive control, the only difference is the fact that the external drive to the two Delay code populations is much weaker, which prevents the circuit from exhibiting double-well dynamics. As there is not persistent activity encoding the upcoming choice, at the Go cue presentation the input from the Delay into the Response neurons is weak and balanced between Left and Right and hence cannot play a role in guiding the licking. Despite the absence of this input, Response populations in response to the Go cue compete to drive a licking response, but the competition in this case is unaffected by the stimulus presented at the beginning of the trial. The tendency to repeat in control-limited phases could be accounted for by an action reinforcement mechanism (Greenstreet et al. 2022; Akaishi et al. 2014; K. J. Miller, Shenhav, and Ludvig 2019) which would either strengthen the synaptic input carrying the Go signal to the Response population driving previous actions (light blue arrow) or by some other mechanism that would facilitate the excitability of the Response neurons that drove those previous actions. We propose that this mechanism could be active in all trials but its impact on biasing the decision dynamics would be overridden by the control signal from the Delay neurons in control-capable trials. The absence of control in some phases would then reveal the presence of this underlying default mechanism.

a

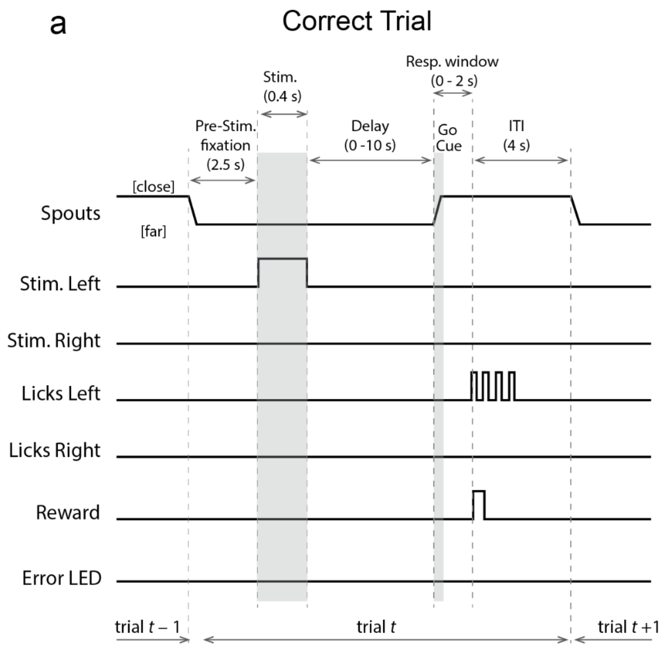


b

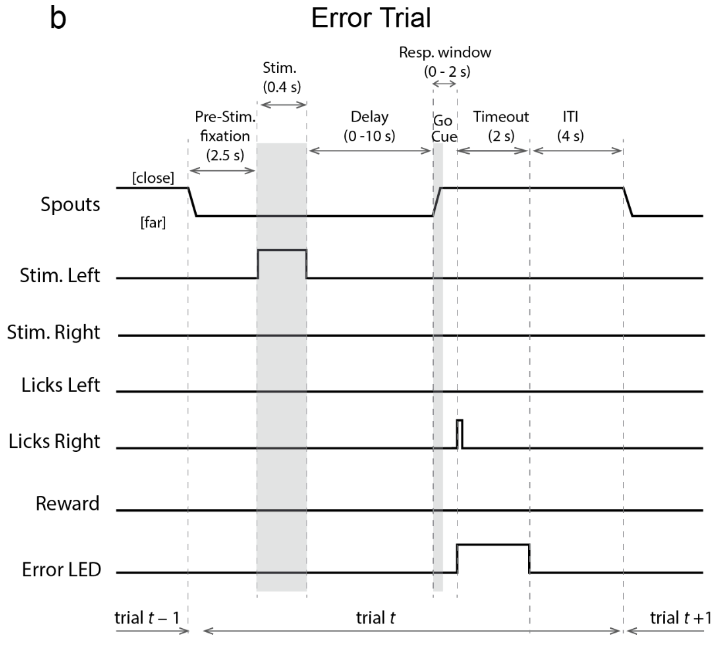


**Supplementary Fig. 12. Time course of the task trial. a,** Time course of a correct Left stimulus trial. **b,** Time course of an incorrect Left stimulus trial. Note that the length of the different trial epochs in the schematic (Stim., Delay, ITI, etc) were not scaled proportionally to their duration. ITI stands for inter-trial-interval.

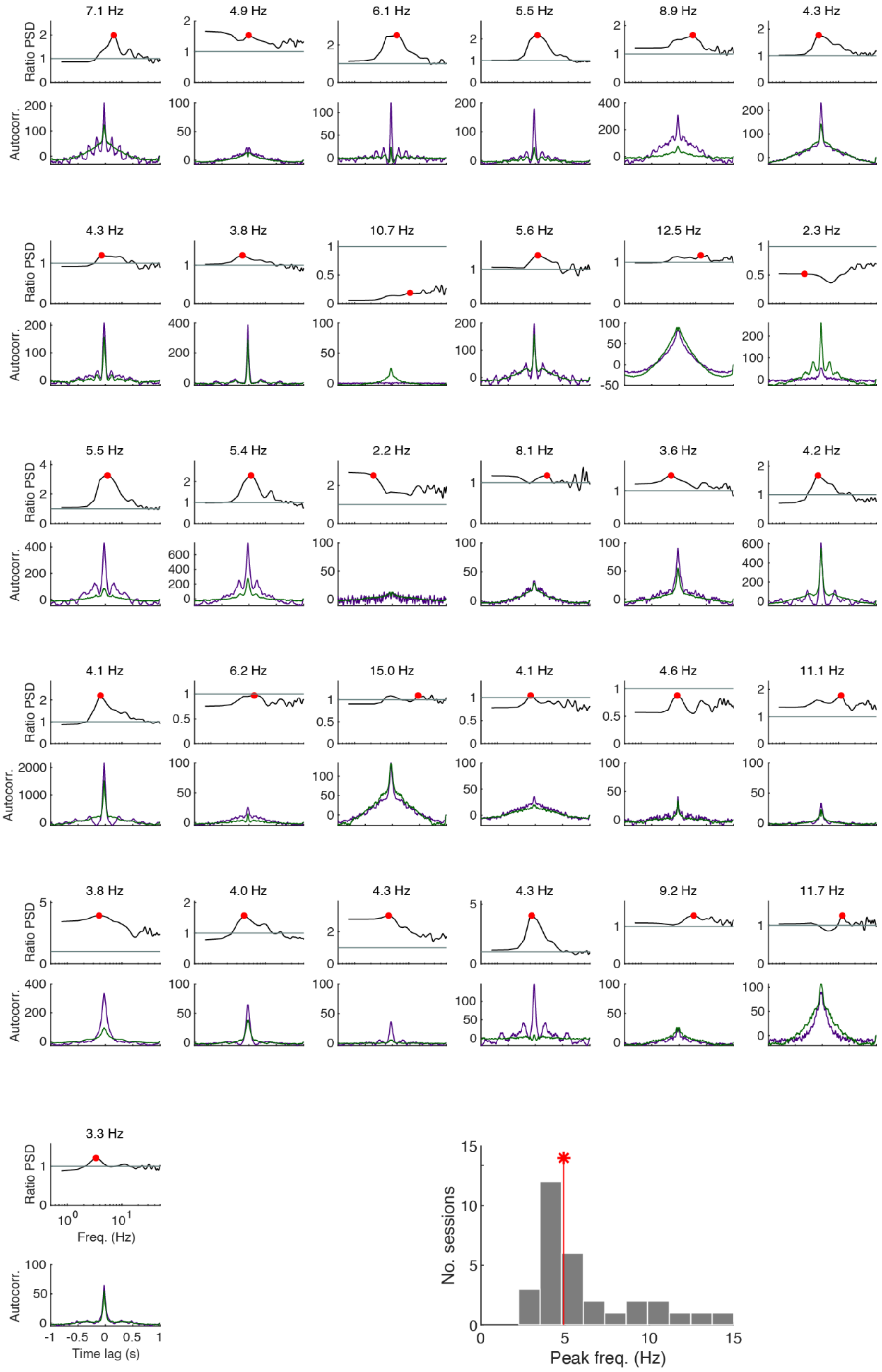


**Supplementary Fig. 13. Population spiking activity in individual sessions shows systematic increase in oscillatory activity during RepL phases.** Every pair of upper-lower panels represents an electrophysiology session (n=31 sessions; Group #3 with n= 8 mice). For each session we show the autocorrelograms (upper panels) and the ratio of power spectral densities (lower panels) computed from the population instantaneous rate during the pre-stimulus period (2 s) in the RepL (purple) and STM (green) phases, as illustrated for the example session shown in Fig. 6e. Red dots show the position of the highest peak in the frequency interval (2, 15) Hz. The horizontal line marks when the ratio equals one for a reference. Bottom: histogram of peak frequencies (horizontal position of red dots) across sessions. Median was 4.91 Hz (red stem) while Q1-Q3 quartiles were 4.06 - 7.88 Hz.
